# Interventionally targeting somatic CAG expansions can be a rapid disease-modifying therapeutic avenue: Preclinical evidence

**DOI:** 10.1101/2025.04.25.650652

**Authors:** Terence Gall-Duncan, Sangyoon Y. Ko, Isabelle K. Quick, Mahreen Khan, Kristie Feng, Chase P. Kelley, Annabelle Coleman, Alexiane Touze, Shuqian Tang, Mustafa Mehkary, Katsuyuki Yokoi, Casey R. Herrington, Justin You, Scott C. Lambie, Tanya K. Prasolava, Gagan B. Panigrahi, Jeehye Park, Kazuhiko Nakatani, Lauren M. Byrne, Peixiang Wang, John S. Schneekloth, Masayuki Nakamori, Paul W. Frankland, Eric T. Wang, Christopher E Pearson

**Author notes:** co-first authors (Terence Gall-Duncan, Sangyoon Y. Ko, Isabelle K. Quick).

## Abstract

Huntington disease (HD) is caused by inherited CAG expansions, which continue expanding somatically in affected brain regions to hasten disease onset and progression. Therapeutically diminishing somatic expansions is expected to be clinically beneficial. However, it is not known if interventionally modifying somatic CAG expansions will actually modify *in vivo* clinically-relevant phenotypes, what the therapeutic window is, or which phenotypes will be altered. Here we show that acute (6-week) delivery of the contraction-inducing slipped-CAG DNA ligand naphthyridine-azaquinolone to young (4-week-old) (CAG)120 HD mice, induces contractions throughout brain regions, improves motor function (locomotion, balance, coordination, muscle strength), molecular disease landmarks (mHTT aggregates, nuclear envelope morphology, nucleocytoplasmic mRNA transport, transcriptomic dysregulation, neuroinflammation), and neurodegeneration. Beneficial effects of modifying somatic expansions were also evident in muscle and blood, where blood CAG instability correlated with brain instability and blood serum had diminished levels of neurofilament light (a biomarker for neurodegeneration) — offering blood as having elements of target engagement and efficacy. These data support that targeting somatic repeat expansions can be a rapid disease-modifying therapeutic avenue for HD and possibly other repeat expansion diseases. Our findings support an etiologic pathway interconnected to somatic CAG expansions that will inform the design of clinical trials expecting clinical benefit by modulating somatic expansions.

## Introduction

Over 15 neurologic, neuromuscular, and neurodegenerative diseases are caused by inherited gene-specific expansions of repetitive CAG DNA sequences, including Huntington disease (HD), several spinocerebellar ataxias (SCAs), and dentatorubral-pallidoluysian atrophy (DRPLA)^1^. While much needed, there are currently no disease-modifying therapies for any repeat disease which would delay, arrest, or reverse the onset or progression of disease^2,3^.

A repeat length threshold for age-dependent disease manifestation is known for the CAG tract in *Huntingtin* (*HTT*)^4^. Specifically, inherited expansions (as measured in blood) of (CAG)≥36-39 manifest with variable low penetrance adult onset, as late as 89 years^5^. Inheritance of (CAG)≥40-55 is fully-penetrant adult onset (presenting at ∼30-65 years). Longer inherited expansions of (CAG)>55 to as many as (CAG)350^6–11^, prompt earlier pediatric or juvenile (JHD) age-of-onset (as early as 12-months), faster progression, and more severe disease^6–8,10–19^. Thus, inherited expansion size drives disease onset, progression, and severity.

Ongoing somatic expansions of inherited CAG lengths occur in affected tissues and neurons with age^20^. For example, HD and JHD brains can show expansions up to (CAG)1000^20–27^. Rates of somatic expansions mirror regional patterns of HD neurodegeneration, with the striatum (vulnerable to degeneration) having the largest expansions and the cerebellum (minimally vulnerable) showing the smallest^26,28–30^. Neurons with larger somatic expansions can exhibit increased vulnerability to cell death relative to cells with stable repeats^28,29,31^. For example, 95– 98% of the highly vulnerable striatal medium spiny neurons (MSNs) from adult-onset HD brains incur expansions of (CAG)35-100 beyond the inherited length, with a smaller portion of MSNs having expansions of (CAG)100-840^28,29,31^. Transcriptomic dysregulation (termed transcriptionopathy) was evident in both populations but was more severe in MSNs with >150 repeats^28,29,31^. This suggests a somatic instability threshold of ∼100 repeats beyond which instability increases and a transition length of >150 repeats where transcriptionopathy worsens^28,29,31^. Thus, somatic expansions in HD correlate with brain region and cell-autonomous pathology.

Targeting somatic CAG expansions is currently one of the most actively pursued disease-modifying therapeutic approaches^2,3^. Interventionally targeting somatic instability has been considered a therapeutic option for >20 years^32–34^. An intense focus on somatic CAG expansions as a therapeutic target arose in 2015 from a series of screens for human modifiers of disease age-of-onset (AOO) or progression in HD individuals, which revealed the DNA repair proteins MSH3, FAN1, PMS1, PMS2, MLH1, and LIG1 as disease modifiers^2,3,35–37^. Disease modifier screens for other repeat expansion diseases including several SCAs, myotonic dystrophy type 1, and X-linked dystonia parkinsonism, have identified the same DNA repair genes^36,38–42^. DNA repair factors identified as AOO modifiers were also known modifiers of somatic expansions, fueling the expectation that modification of somatic expansions would be an attractive disease-modifying therapeutic target^43^. Mechanistically, spontaneous CAG expansions in HD mice require MSH3, while FAN1 suppresses hyper-expansions. MSH3 and FAN1 are thought to form and process slipped-CAG DNA structures, mutagenic intermediates of expansion mutations^28^. Slipped-CAG structures are a critical element in all models of instability^28,31,44–49^. To this degree targeting slipped-CAGs, inhibiting MSH3 or overexpressing FAN1 might be expected to modify somatic expansions and disease. These studies are based upon aged animals born from pre-zygotic genetic ablation of CAG instability modifying genes.

In contrast to animal models in which repeat-modifying genes can be knocked-out pre-zygotically, HD patients would be treated postnatally with an anti-CAG expansion therapy, were one available. Studies of active expansions in premanifest *HTT* expansion carriers suggests an early but wide therapeutic window by which interventional modulation of expansions could provide clinical benefit^31,47^. For example, in *HTT* expansion carriers blood the levels of CAG expansions and neurofilament light (NfL; a biomarker of neurodegeneration) increase in parallel to striatal atrophy over decades prior to disease onset^47^. Thus, strong correlative data supports early targeting of somatic CAG expansions as a disease-modifying therapeutic approach.

To date there is no demonstration of clinical benefit following interventional targeting of somatic CAG expansions. Preclinical evidence of *in vivo* benefit of HD-relevant phenotypes following interventional modulation of somatic CAG expansions will provide validation of the therapeutic target. Moreover, such *in vivo* evidence-based awareness can provide guidance for clinical trial design.

Five critical questions remain unanswered regarding targeting somatic CAG expansions as a disease-modifying therapeutic approach: First *efficacy*, can interventional modulation of somatic expansions actually lead to benefit of clinically-relevant phenotypes in an *in vivo* model? Second, what phenotypes (motor, molecular, and cellular disease landmarks) might be detectably affected by interventional modulation of somatic repeat expansions? Third, what is the *therapeutic window* during which administration of a possible intervention might lead to detectable improvements of clinically-relevant phenotypes. Fourth, what is the *time-to-clinical benefit* that might be hoped for when interventionally modifying somatic expansions? Fifth, can interventional modulation of somatic expansions and their benefits be monitored in a readily accessible tissue?

Here, with the goal of testing whether interventional modulation of somatic CAG expansions can be a disease-modifying therapeutic approach to benefit HD-related phenotypes, we used the small molecule naphthyridine-azaquinolone (NA), previously demonstrated to induce *en masse* contractions of the mutant CAG-expanded allele to less than inherited lengths in brains of HD or DRPLA mice^49,50^.

NA targets active somatic expansions with a known mechanism of action. NA binds only to slipped-CAG structures, which only occur in the mutant repeat during active expansion mutations^49^. Due to this exquisite specificity, NA did not induce off-target mutagenesis as assessed via whole genome sequencing, deep-long read sequencing of *HGPRT* (a mutation load proxy), cancer MSI panel, and repeat sizing of non-expanded CAG-containing genes. Moreover, NA was not toxic, tolerable in mice for at least 22-weeks with no adverse events or pathology, and did not affect transcription or translation of wildtype or mutant *HTT* or *ATN1* genes^49,50^. The mechanism by which NA-mediates CAG contractions, as with spontaneous expansions, requires transcription across the expanded repeat, MSH3, and FAN1^49,51^. These proof-of-principle studies highlight NA’s utility to address whether interventionally modulating somatic expansions might benefit HD-related phenotypes.

## Results

### Inhibited somatic expansions, induced contractions, and diminished polyQ aggregates in multiple brain regions

To assess the effects of NA on HD phenotypes, we administered NA for 6-weeks into brains of R6/2 mice confirmed to inherit 120-121 CAG repeats (determined in tail DNA at 2-weeks of age) (Fig 1A). We opted to use R6/2 mice, one of the most extensively characterized HD models with known timings of HD-related phenotypes (Extended Fig S1A). R6/2 mice show aggressively progressing, age-dependent, and tissue-specific somatic CAG expansions, aberrant neurodevelopment, polyQ aggregates, transcriptionopathy, neuroinflammation, microglial activation, cumulatively increasing levels of NfL, motor defects, and a shortened lifespan (∼13-15 weeks)^52–73^. Striatal somatic CAG expansions are just evident at 4-weeks of age in mice devoid of symptoms and ramp-up over 6-10-weeks, coincident with the onset of overt motor symptoms^20,21,26,52,74^. Similar to humans, R6/2 mice show somatic expansions, aggregates, and elevated NfL prior to overt disease^52,53,62–64,69,70^. Awareness of these timings of molecular, pathological, and motor defects served to guide our dosing regimen (Extended Fig S1A).

**Figure 1.**
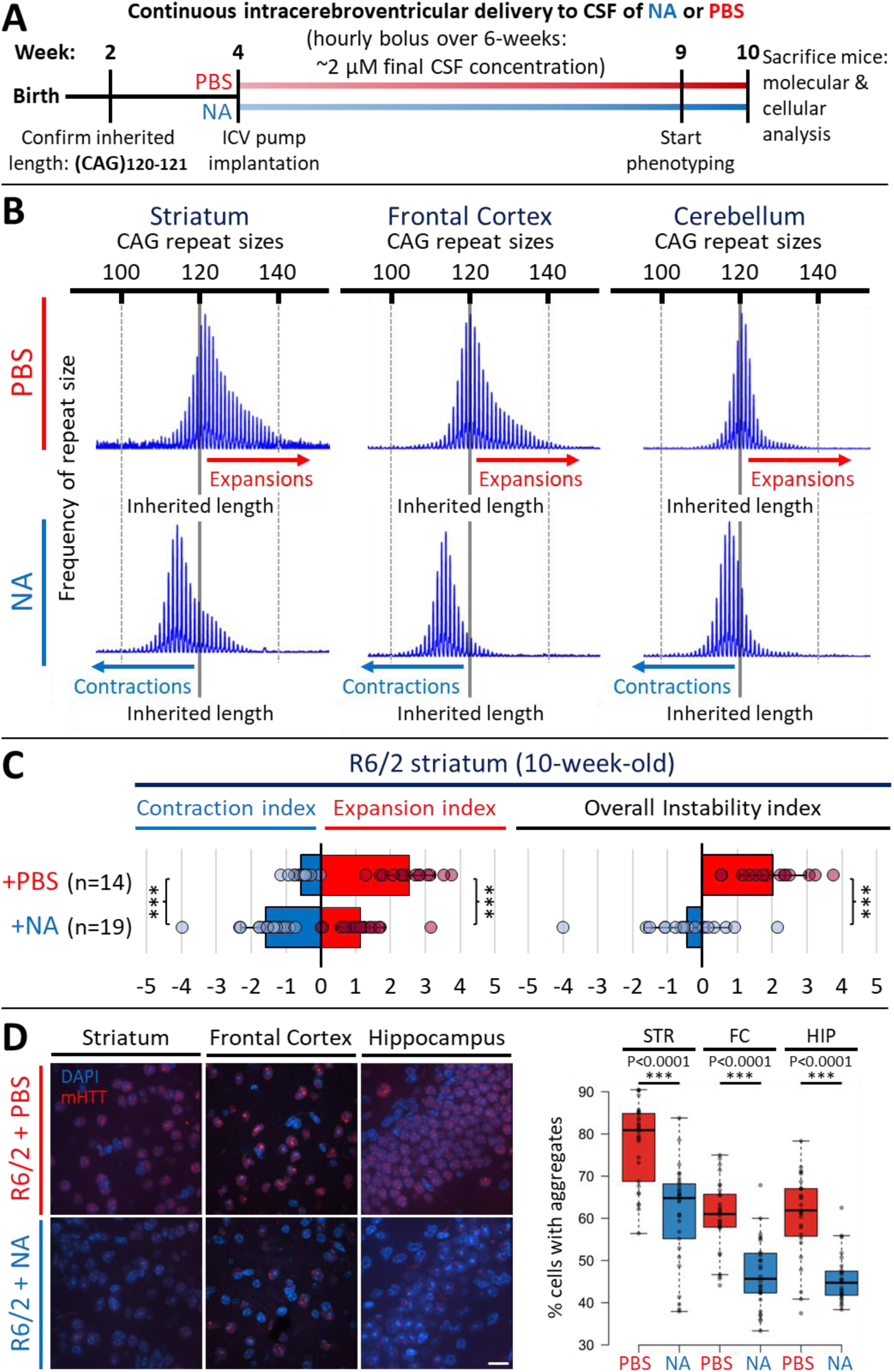
ICV-delivered NA prevents expansions, induces *en masse* contractions, and diminishes aggregates in multiple R6/2 HD mouse brain regions. **A)** Summary of treatment regimen, treatment timeline, phenotypic assessments, and tissue harvesting for mice used in this study. **B)** Representative fragment length analysis (FLA) scans from Peak Scanner software (ThermoFisher Scientific) outlining repeat instability in brain regions of 10-week-old R6/2 mice +/−NA. Inherited length (CAG)_120-121_ is denoted with a solid gray line, additional reference lengths (+/− 20 CAG units from the inherited (CAG)_120-121_) are denoted using dashed gray lines. Expansion-biased peaks are denoted with a red arrow while contraction-biased peaks are denoted with a blue arrow. **C)** Striatal repeat expansion indices denoting the average expansion/contraction indices (left panel) and overall instability indices (right panel) in 10-week-old R6/2 mice treated with vehicle (n = 14) or NA (n = 19). Bars denote average instability indices, error bars represent standard deviation from the mean, and circles represent values for individual mice. Statistics: one-tailed Mann-Whitney U test, *** denotes p<0.0001. **D)** Left panel: representative 40x confocal images outlining mutant poly-glutamine aggregates in the striatum, frontal cortex and hippocampus of 10-week-old R6/2 mice +/−NA. Red signal denotes mutant HTT (mHTT) poly-glutamine fragment aggregates, blue signal denotes DAPI-stained nuclei. Scale bar = 16 μm. Right panel: quantification of percentage of cells with nuclear poly-glutamine aggregates relative to all nuclei per 40x image in the striatum, frontal cortex, and hippocampus of 10-week-old R6/2 mice +/−NA (n = 10 images per mouse, ∼300-500 cells assessed total per mouse, with n = 3 mice per condition). Box denotes 1^st^ to 3^rd^ quartiles, black lines denote median, Tukey whisker extents. Statistics: unpaired t-test, *** denotes p<0.0001.

NA or PBS (vehicle control) was continuously delivered to the cerebrospinal fluid (CSF) intracerebroventricularly (ICV) by an ALZET® osmotic pump subcutaneously implanted into 4-week-old mice, when somatic expansions begin. Implanted pumps allow for untethered spontaneous movement, reducing the need for invasive handling during delivery. An hourly bolus of 0.15 μl/hr of a 500 μM NA stock yielded an estimated ∼2 μM in the CSF per bolus, assuming a CSF volume of ∼36 microliters per mouse^75,76^. Cultured HD patient fibroblasts show a lack of toxicity up to 50 μM, and previous ICV delivery of 8 μM NA for 22-weeks in DRPLA mouse brains was non-toxic and tolerable, supporting that ∼2 μM CSF concentration used here is also safe, non-toxic and tolerable^49,50^. After 5-weeks of NA delivery, mice were evaluated for locomotion, grip strength, motor coordination and clasping deficits. Mice were continuously treated during evaluations and were sacrificed at 10-weeks of age for tissue harvesting (after 6-weeks of NA administration) (Fig 1A; Extended Fig S1A). A summary of findings for individual mouse can be found in Extended Table S1.

NA induced CAG contractions in multiple brain regions, indicating its ability to distribute through the brain via the CSF. Vehicle-treated mice (n = 14) incurred maximal spontaneous expansions of up to ∼140, ∼140, and ∼128 repeats above the inherited (CAG)120 in the striatum, frontal cortex, and cerebellum, respectively, over the course of the experiment (Fig 1B-C). NA-treated mice (n = 19) incurred significantly inhibited expansions and induced contractions below the inherited length relative to vehicle-treated R6/2 mice; within the striatum and frontal cortex, average contractions of 6-8 repeat CAG units below inherited size were observed for most NA-treated mice, and maximum repeat sizes were on average 10-20 CAG units smaller than the maximum size observed in vehicle-treated R6/2 mice (Fig 1B).

Quantitatively, average instability indices of +2 (expansion-bias) were observed in vehicle-treated R6/2 striata compared to −0.4 (contraction bias) in NA-treated R6/2 striata (Fig 1B-C, p < 0.00001 for instability, contraction, and expansion indices). Similarly, average instability indices of +1.2 were observed in vehicle-treated R6/2 frontal cortex compared to −0.6 in NA-treated R6/2 frontal cortex (Fig 1B, Extended Fig S1B, p < 0.00001 for instability and expansion indices, p = 0.0002 for contraction index). Relative to vehicle-treated R6/2 cerebellum (instability index: +0.3), NA-treated R6/2 cerebellum exhibited milder, yet significant, inhibition of expansions (p = 0.05) and NA-induced contractions of 2-3 repeat units on average (p = 0.003) with an overall instability index of −0.7 (Fig 1B and Extended Fig S1B; p = 0.007 for instability index). Overall levels of NA-induced contractions were greater in brain regions with greater spontaneous expansions in vehicle-treated mice (Fig 1B).

Coincident with contracted repeats, NA diminished toxic polyQ aggregates (Fig 1D)^77–80^. NA significantly reduced the percentage of neurons expressing aggregates within the striatum, frontal cortex, and hippocampus (Fig 1D, aggregates mouse quantifications, Extended Fig S1C, individual mouse quantifications). NA did not significantly alter aggregates within cerebellar granular cells or Purkinje neurons (Extended Fig S1B-C), consistent with NA’s limited effect on cerebellar CAG instability (Fig 1B, Extended Fig S1B).

These data extend our previous studies of striatally-targeted delivery of NA^49^, and now reveal that NA delivered to the CSF can disseminate to multiple brain regions to inhibit expansions, induce contractions, and reduce polyQ aggregate formation.

### Induced CAG contractions improved motor phenotypes

Progressive loss of motor functions is a prominent and cardinal disease symptom in HD patients and mouse models^60^. We assessed NA modulated motor phenotypes using four tests on 9-10-week-old WT or R6/2 mice following 5-6 weeks delivery of NA or vehicle: 1) open field test for general locomotion; 2) hanging wire test for grip strength; 3) accelerating rotarod for both motor coordination and learning; and 4) hindlimb clasping for cerebellum-associated motor deficits. Having both NA- and vehicle- treated groups of WT mice allowed for the assessment of any non-specific effects of NA in healthy control mice.

NA rescued locomotor activity impairment in open field tests (Fig. 2A and 2B). WT mice treated with NA (n = 7) or vehicle (n = 10) exhibited a comparable total distance travelled, indicating that NA did not have non-specific effects on general locomotion in healthy control mice (Fig 2A, quantification, and Fig 2B, aggregate heatmaps). Vehicle-treated R6/2 mice (n = 12) exhibited significantly reduced total distance traveled compared to both WT groups, traveling ∼1/2 the distance on average (Fig 2A and 2B, p < 0.0001), confirming HD-like motor deficits. In contrast, NA-treated R6/2 mice (n = 16) showed significantly increased total distance traveled compared to vehicle-treated R6/2 mice (Fig 2A and 2B, p = 0.0317). Although NA enhanced locomotor activity in R6/2 mice, overall activity still remained lower than WT mice (Fig 2A and 2B, p < 0.0001). These data suggest that NA partially alleviated locomotor deficits in R6/2 mice.

**Figure 2.**
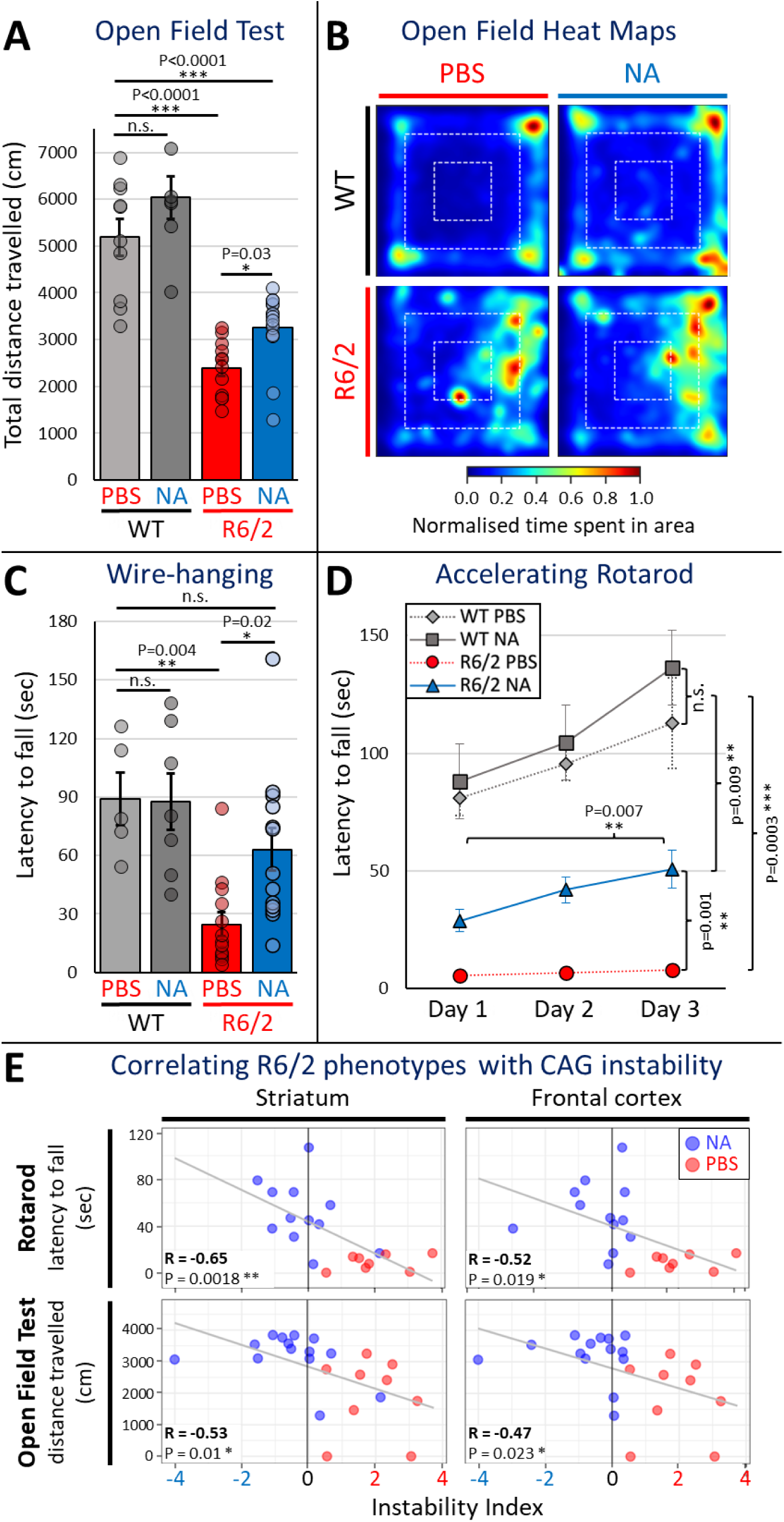
NA improves motor phenotypes in R6/2 HD mice. **A)** Reduced locomotor activity in R6/2-PBS mice is recovered in R6/2-NA mice during open field test (two-way ANOVA; Treatment effect: *F*_1,41_ = 9.14, *P* = 0.0043; *n* = 10 (WT-PBS), 7 (WT-NA), 12 (R6/2-PBS), 16 (R6/2-NA)). **B)** Grouped heatmaps outlining increased center zone time in vehicle-treated R6/2 mice due to reduced locomotor activity, whereas R6/2-NA mice exhibit improved movement patterns similar to WT-PBS and WT-NA mice (*n* = 10 (WT-PBS), 7 (WT-NA), 12 (R6/2-PBS), 16 (R6/2-NA)). White dashed lines indicate the boundaries of the center, middle, and outer zones. **C)** R6/2-PBS mice exhibit reduced latency to fall in the wire-hanging test, whereas R6/2-NA mice recover to the typical level observed in WT-PBS and WT-NA mice (two-way ANOVA; Genotype × Treatment interaction: *F*_1,34_ = 2.94, *P* = 0.0957; *n* = 5 (WT-PBS), 7 (WT-NA), 13 (R6/2-PBS), 13 (R6/2-NA)). **D)** R6/2-PBS mice exhibit reduced latency to fall compared to WT-PBS and WT-NA mice across a 3-day rotarod motor learning task, whereas R6/2-NA mice show improved latency to fall (three-way repeated measures ANOVA; Genotype × Day interaction: *F*_2,70_ = 7.66, *P* = 0.0010, Treatment × Day interaction: *F*_2,70_ = 3.03, *P* = 0.0546, Treatment effect: *F*_1,35_ = 5.34, *P* = 0.0269; *n* = 11 (WT-PBS), 6 (WT-NA), 10 (R6/2-PBS), 12 (R6/2-NA)). Phenotypic data in A-D are shown as mean ± SEM. \**P* < 0.05, \*\**P* < 0.01, \*\*\**P* < 0.001, \*\*\*\**P* < 0.0001, n.s., not significant. **E)** Pearson correlations comparing striatal instability indices (left panels) or frontal cortex instability indices (right panels) to rotarod latency to fall (top panels) or distance travelled in an open field (bottom panels), ** denotes p<0.01 and * denotes p<0.05.

NA rescued impairments in grip strength in the hanging wire test (Fig. 2C). WT mice treated with NA (n = 7) or vehicle (n = 5) did not show changes in latency to fall from the hanging wire, again confirming a lack of non-specific effects in healthy control mice (Fig 2C). Vehicle-treated R6/2 mice (n = 13) showed significant reduction in latency compared to both WT groups, with roughly one-third of the latency to fall on average (Fig 2C, p < 0.005), consistent with reduced grip strength in R6/2 mice. NA-treated R6/2 mice (n = 13) exhibited significantly longer latencies to fall compared to vehicle-treated R6/2 mice (Fig 2C, p = 0.0222). Additionally, latency in NA-treated R6/2 mice are comparable to that of both WT groups (Fig 2C, p = 0.3274), suggesting that NA effectively rescued grip strength deficits in R6/2 HD mice to WT levels.

NA treatment rescued impairments of motor coordination and learning in accelerated rotarod testing (Fig. 2D). Motor coordination was assessed by measuring latency to fall from the rotarod, with daily trials conducted over 3 days. WT mice treated with NA (n = 6) or vehicle (n = 11) did not exhibit differences in latency to fall from the rotarod (Fig 2D), again confirming a lack of non-specific effects in healthy control mice. Vehicle-treated R6/2 mice (n = 10) exhibited significant reductions in latency to fall, remaining on the rod for ∼1/20^th^ of the time compared to both WT groups on the third day of testing (Fig 2D, p < 0.005), indicating severe motor coordination deficits in R6/2 mice. In contrast, NA-treated R6/2 mice (n = 12) demonstrated robust improvements on the third day of testing, with a ∼10-fold increase in latency to fall compared to vehicle-treated R6/2 mice (Fig 2D, p = 0.0013), though they still showed ∼1/2 the latency of both WT groups (Fig 2D, p = 0.0095).

Rotarod data was also used to assess for motor learning by analyzing day-to-day improvements in latency to fall within each group over the 3 days of testing. Both WT groups showed significant increases in latency (Fig 2D, p < 0.05), indicating motor skill acquisition which was unaffected by NA in healthy control mice. Vehicle-treated R6/2 mice did not exhibit such improvement (Fig 2D), suggesting that R6/2 mice are not acquiring motor skills (i.e. learning) from their daily trials. In contrast, NA-treated R6/2 mice exhibited significant improvement between day 1 and day 3 of testing (Fig 2D, p = 0.0068). These data suggest that NA-induced CAG contractions alleviated deficits in motor coordination and learning in R6/2 HD mice.

NA did not rescue cerebellar ataxia as measured by hindlimb clasping (Extended Fig S2A). WT mice treated with NA (n = 7) or vehicle (n = 6) showed no differences in average clasping scores (Extended Fig S2A), demonstrating an absence of non-specific effects. Vehicle-treated R6/2 mice (n = 14) exhibited a significant increase in average clasping scores compared to both WT groups (p < 0.005), indicating ataxia in R6/2 mice. NA-treated R6/2 mice (n = 14) did not show significant improvements in hindlimb clasping (p = 0.0862). The absence of an effect of NA in this test is consistent with the limited effect of NA upon either CAG instability and polyQ aggregates within the cerebellum (Extended Fig S2A), as hindlimb clasping is primarily associated with cerebellar dysfunction^81,82^.

### Levels of interventionally-modified expansions correlate with improved motor phenotypes

To assess if somatic instability is associated with disease phenotypes, levels of somatic instability within each brain region were correlated with phenotypic scores. Overall somatic instability indices (sum of individual expansion and contraction indices) in the striatum negatively correlated with latency to fall from an accelerating rotarod (Fig 2E, R = −0.65, p = 0.0018) and distance travelled in an open field (Fig 2E, R = −0.53, p = 0.01); in the frontal cortex, instability negatively correlates with latency to fall from an accelerating rotarod (Fig 2E, R = −0.52, p = 0.019) and distance travelled in an open field (Fig 2E, R = −0.47, p = 0.023). Latency to fall from a hanging wire and hindlimb clasping scores did not correlate with instability in any tissue, and levels of instability in the cerebellum did not correlate with any phenotypic scores (Extended Fig S2B-C). Cumulatively, these data support that interventionally reduced levels of somatic CAG expansions in the striatum and frontal cortex correlate with better open field and rotarod scores.

To assess whether phenotypic rescue of open field test and rotarod scores depend on contractions or expansions (or both) within specific brain regions, parsed contraction and expansion indices were separately correlated with individual phenotypic domains. Open field test scores significantly correlated with striatal and frontal cortex expansion indices. (Extended Fig S2B). In contrast, rotarod performance significantly correlated with contraction indices in the striatum, as well as expansion indices in the striatum and frontal cortex (Extended Fig 2C).

Together, these data support that NA-mediated inhibition of expansions in the striatum and frontal cortex play a larger role in open field test performance (locomotor activity) relative to contractions. In contrast, both NA-induced contractions in the striatum and inhibition of expansions in the striatum and frontal cortex contribute to improved rotarod performance (motor coordination). As such, brain-region specific patterns of contractions and expansions differentially correlate with specific phenotypic presentations.

### Induced CAG contractions improved nuclear envelope morphology and nuclear mRNA retention in neurons

Since NA diminished toxic polyQ aggregate levels in R6/2 striata (Fig 1D and Extended Fig S1B), we next assessed if downstream aggregate-associated pathogenesis were also rescued. Aggregates in HD patient and R6/2 mouse brains accelerate ageing in MSNs, in part by sequestering nuclear pore complex proteins, which induces abnormal age-related nuclear envelope morphology, which disrupts nuclear functions^83–87^. In neurons of young unaffected individuals, the nuclear membrane, as assessed by Lamin B1 (a major component of the nuclear envelope), appears spherically smooth, typical of a normal young nuclear envelope. In contrast, in aged unaffected individuals and young HD patients, the nuclear membrane becomes aged with invaginations and wrinkles. We assessed the effect of interventionally-modified CAG expansions upon nuclear envelope morphology of striatal cells.

Vehicle-treated R6/2 mice showed significantly more (∼43%) MSNs with aberrant nuclear membranes relative to age-matched vehicle-treated R6/2 mice (∼25%), consistent with accelerated aging in HD patients and mice (Fig 3A, p < 0.0001, representative images and aggregate mouse quantifications, and Extended Fig S3A, individual mouse quantifications)^88^. In contrast, NA-treated R6/2 striata exhibited significantly fewer neurons (∼30%) with abnormal nuclear envelope morphology relative to vehicle-treated R6/2 mice (Fig 3A p < 0.0001, and Extended Fig 3A). NA-treated R6/2 mice still exhibit significantly elevated abnormal nuclear envelope morphology relative to WT mice, suggesting partial rescue (Fig 3A p = 0.005, Extended Fig S3A).

**Figure 3.**
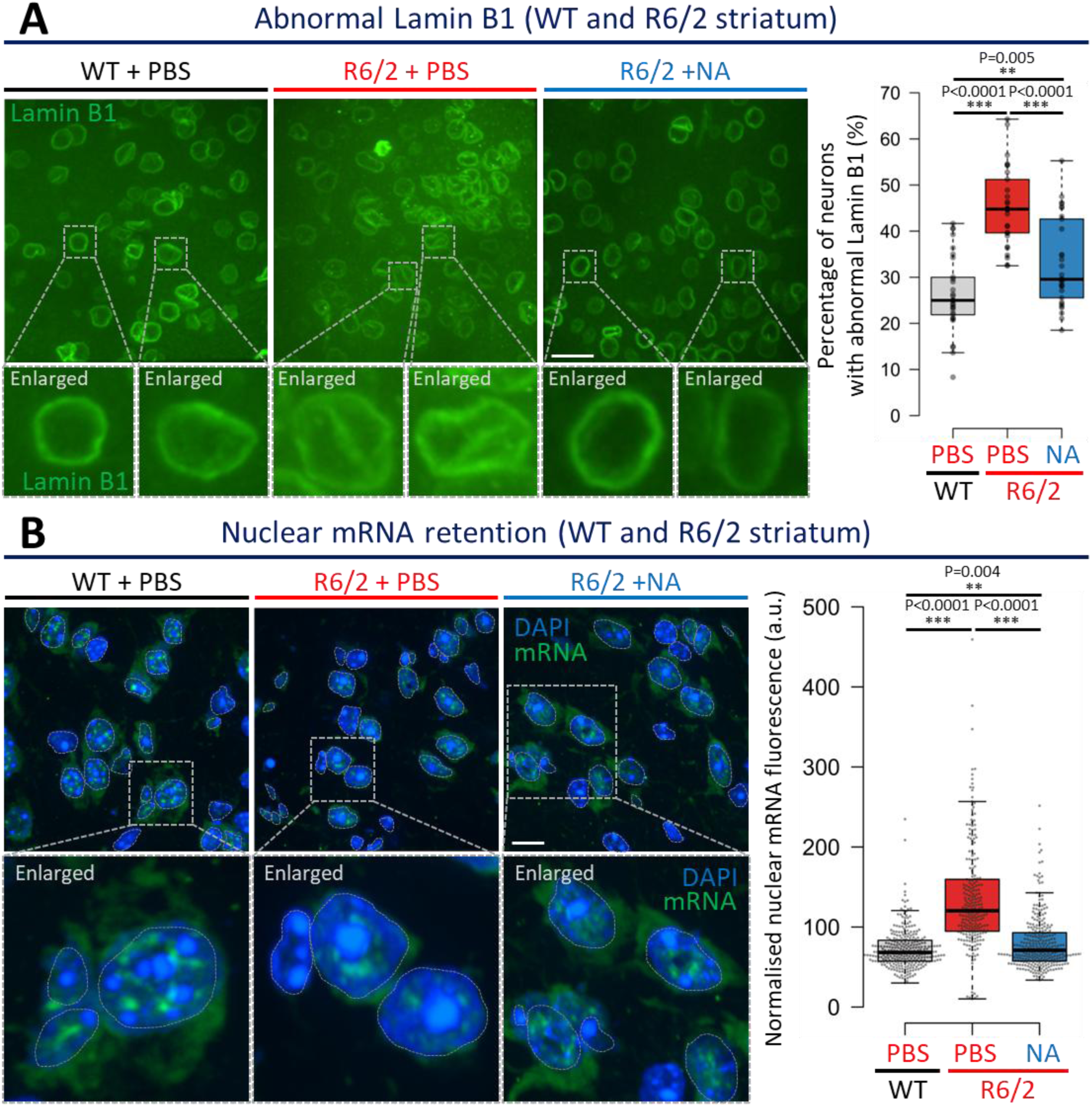
NA improves nuclear envelope morphology and mRNA nuclear export in R6/2 HD striata. **A)** Left panels: representative 40x confocal images outlining Lamin B1 in striatal cells of 10-week-old R6/2 mice +/−NA. Green signal denotes Lamin B1 (nuclear envelope marker), blue DAPI signal was removed in representative images to increase clarity of Lamin B1 signal visualisation. Scale bar = 16 μm. Dotted boxes and lines denote enlarged individual cells for visualisation purposes. Right panel: quantification of percentage of cells with abnormal Lamin B1 morphology relative to all nuclei per 40x image in the striatum of 10-week-old R6/2 mice +/−NA (n = 10 images per mouse per condition, ∼350-550 cells assessed total per mouse, with n = 3 mice per condition). Box denotes 1^st^ to 3^rd^ quartiles, black lines denote median, Tukey whisker extents. Statistics: unpaired t-test, *** denotes p<0.0001. **B)** Left panels: representative 40x confocal images outlining mRNA retention in striatal cells of 10-week-old R6/2 mice +/−NA. Green signal denotes mRNA (polyT RNA FISH), blue signal denotes DAPI-stained nuclei, dotted grey line overlay denote the nuclei boundary for visualisation purposes. Scale bar = 16 μm. Dotted boxes and lines denote enlarged individual cells for visualisation purposes. Right panel: quantification of normalised nuclear mRNA fluorescence for cells in the striatum of 10-week-old R6/2 mice +/−NA (n = ∼125 individual cells assessed over 2 technical replicates per mouse, each replicate assessing all nuclei within 8-10 40x images, n = 3 WT mice and NA-treated R6/2 mice and n = 2 PBS-treated R6/2 mice). Box denotes 1^st^ to 3^rd^ quartiles, black lines denote median, Tukey whisker extents. Statistics: unpaired t-test, *** denotes p<0.0001, ** denotes p<0.01.

Impeded mRNA transport out of the nucleus is a functional consequence of abnormal nuclear envelope morphology, as previously observed in HD patient and HD mouse brains^84^. mRNA retention could contribute to wide-spread transcriptionopathy – a core pathological feature of HD which drives neuronal death. Vehicle-treated R6/2 mice striatal neurons exhibited mRNA signal localised almost entirely within their nuclei, demonstrating significantly higher mRNA nuclear retention than vehicle-treated WT striata (Fig 3B p < 0.0001 and Extended Fig S3B). In contrast, NA-treated R6/2 striatal neurons mRNA retention was significantly reduced relative to vehicle-treated R6/2 striata (p < 0.0001) (Fig 3B). In both NA-treated R6/2 and vehicle-treated WT striatal neurons, mRNA is observed both inside and outside of the nucleus, consistent with effective mRNA transport out of the nucleus (Fig 3B, representative images and aggregate mouse quantifications, and Extended Fig S3B, individual mouse quantifications). Rescue of mRNA retention by NA was partial as retention was significantly elevated relative to WT mice (p = 0.004) (Fig 3B and Extended Fig S3B).

### Induced CAG contractions improved transcriptionopathy

Transcriptionopathy is evident in HD patient brains, is an early event in HD mice (6-weeks in R6/2), is exacerbated by both age and longer CAG lengths^65,89^, and dysregulated gene sets are similar between human HD patients relative to full-length and N-terminal HTT-fragment HD mice (albeit temporally distinct)^55,61,65,90,89,91,92^. To assess if NA treatment affected transcriptionopathy, we conducted RNA-seq on striata of NA- or vehicle-treated R6/2 mice relative to WT mice (n = 3/group; Extended Table 2). Differentially expressed genes (DEGs) were defined as having a +/− 0.5 log2FC value and a false-discovery rate (FDR)-adjusted p < 0.05.

Vehicle-treated R6/2 striata exhibited 1216 DEGs, 837 down-regulated and 379 up-regulated relative to WT striata (Fig 4A). NA-treated R6/2 striata exhibit 27% fewer dysregulated genes, with 687 down-regulated and 201 up-regulated genes relative to WT striata (Figure 4B). Up- regulated genes mostly overlapped between vehicle- and NA-treated R6/2 striata, although 47 are up-regulated in NA- but not in vehicle-treated R6/2 striata (Fig 4C). Down-regulated genes overlapped less, with 311 genes down-regulated in NA- but not in vehicle-treated R6/2 striata (Fig 4C). Together, these 358 DEGs (47+311) represent unique changes in R6/2 striata due to NA, though this is likely an overestimate due to limited statistical power from sample size (n = 3/condition). Conversely, there are 686 DEGs (225+461) in vehicle- but not in NA-treated R6/2 striata relative to WT, representing DEGs in R6/2 whose expression is likely rescued by NA towards WT levels, with similar statistical limitations.

**Figure 4.**
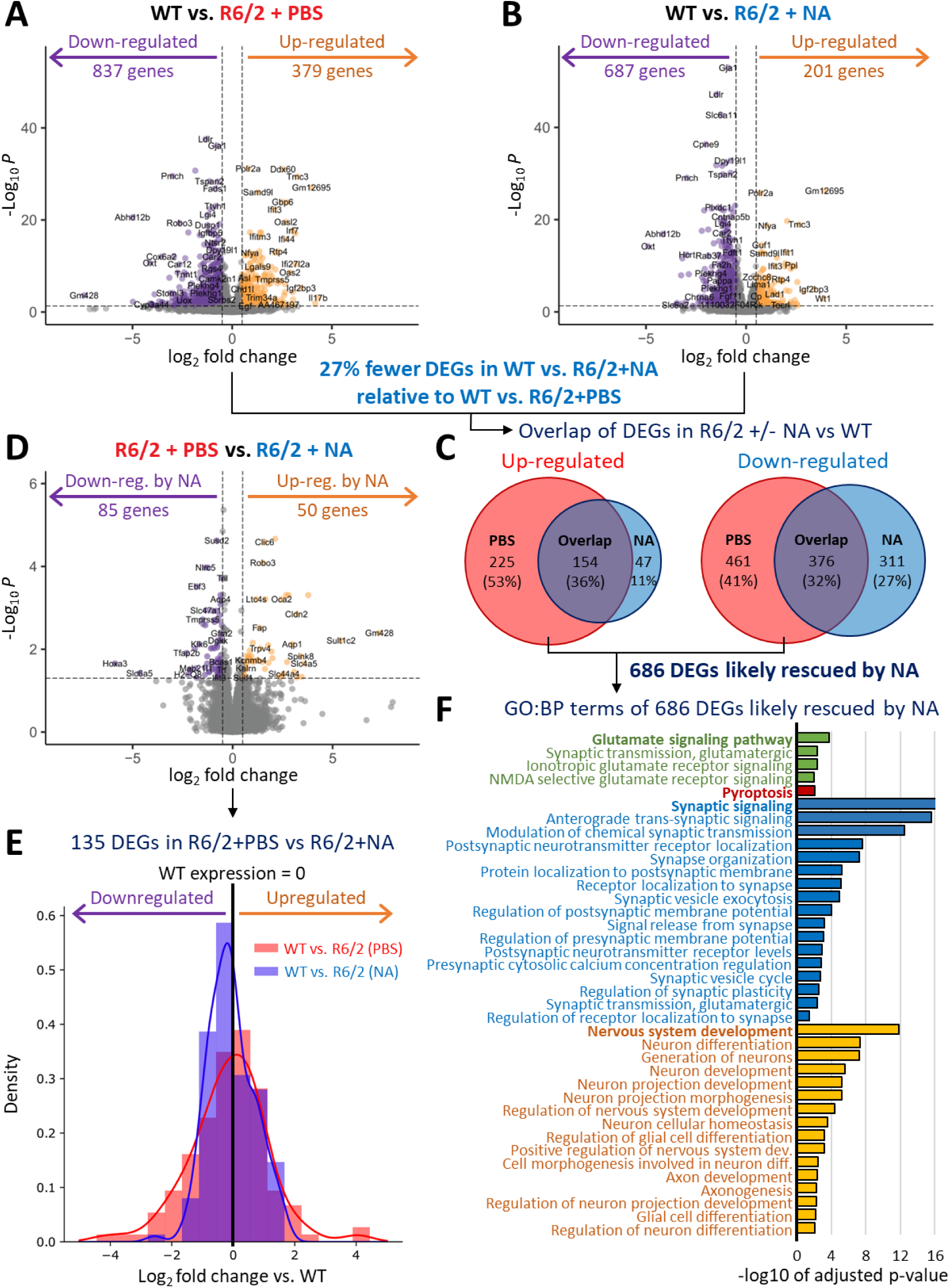
NA mitigates disease-associated transcriptionopathy in R6/2 HD striata. A-B) Volcano plot comparing differentially expressed genes (DEGs) in 10-week-old A) vehicle-treated R6/2 striata relative to vehicle-treated WT striata and B) NA-treated R6/2 striata relative to vehicle-treated WT striata (n = 3 mice per condition). X-axis dotted lines correspond to +/−0.5 log2FC and the Y-axis dotted line corresponds to an FDR-adjusted p<0.05, where dysregulated genes are defined as having a +/−0.5 log2FC value and a false-discovery rate (FDR)-adjusted p < 0.05. Purple dots represent downregulated genes while orange dots represent upregulated genes, relative to WT. **C)** Venn Diagrams outlining the number of unique and overlapping upregulated (left) and downregulated (right) DEGs in R6/2 striata +/−NA vs vehicle-treated WT striata. **D)** Volcano plot comparing differentially expressed genes (DEGs) in vehicle-treated R6/2 striata relative to NA-treated R6/2 striata relative to vehicle-treated WT striata (n = 3 mice per condition). X-axis dotted lines correspond to +/−0.5 log2FC and the Y-axis dotted line corresponds to an FDR-adjusted p<0.05, where dysregulated genes are defined as having a +/−0.5 log2FC value and a false-discovery rate (FDR)-adjusted p < 0.05. Purple dots represent downregulated genes while orange dots represent upregulated genes, relative to WT. **E)** histogram outlining the density of gene expression in R6/2 striata +/−NA as a function of log2fold change relative to WT for 135 DEGs identified in vehicle treated R6/2 striata vs NA-treated R6/2 striata. Red bars and associated trend line denote expression of these genes within the vehicle-treated R6/2 striata, while blue bars and associated trend line denote NA-treated R6/2 striata. WT expression is denoted by the middle black line connotating log2foldchange=0. **F)** Gene ontology: biological function (GO:BP) terms for 686 DEGs which are likely rescued by NA, with the x-axis denoting the −log10 of adjusted p-value. Bolded terms are the 4 main terms associated with subsequent sub-terms; glutamate signalling associated with green terms, pyroptosis associated with red terms, synaptic signalling associated with blue terms, and nervous system development associated with yellow terms.

Vehicle-treated R6/2 striata expression was also directly compared to NA-treated R6/2 striata (Figure 4D). NA-treated R6/2 striata exhibit 135 DEGs relative to vehicle-treated R6/2 striata - 85 genes down-regulated and 50 genes up-regulated (Figure 4D). Comparing the expression of these 135 genes to WT mice, NA-treated R6/2 striata expression was more similar to WT striata while vehicle-treated R6/2 striata deviated more strongly from WT (Fig 4E; compare NA expression in blue to vehicle expression in red). This is consistent with the hypothesis that NA treatment reduces transcriptionopathy in R6/2 striata.

### Induced CAG contractions mitigated expression in HD-related pathways

To assess if NA-mediated transcriptional changes were relevant to HD-related pathways, we evaluated the gene ontology (GO) terms associated with 1) the 686 NA-rescued DEGs, and 2) the 358 DEGs uniquely elicited by NA in R6/2 striata relative to WT (Extended Table 3).

The 686 NA-rescued DEGs were enriched in GO pathways known to be dysregulated in HD patient brains (Fig 4F, Extended Table 3). Four major paths include: 1) glutamate signaling, where polyglutamine-prompted excitotoxicity drives neuron dysfunction/death^93–96^, 2) pyroptosis, where neuroinflammatory responses promote neurodegeneration^66,97–99^, 3) synaptic signaling, where neuron-to-neuron communication is lost within and between brain regions^93, 100–103^, and 4) nervous system development^104–108^.

Expression levels of some of the 686 NA-mitigated DEGs in each of the mouse groups from the 4 pathways outlined above are visualized by heatmaps in Extended Fig 4A. Glutamate signaling pathway genes were down-regulated in vehicle-treated R6/2 striata, consistent with loss of glutamate signaling in HD patients^93–96^, with NA mitigating their expression. Most inflammation/pyroptosis genes were up-regulated in vehicle-treated R6/2 striata, consistent with up-regulation of neuroinflammation in HD patients^105–108^, and down-regulated by NA. Synaptic signaling genes were predominantly down-regulated in vehicle-treated R6/2 striata, consistent with a loss of neuron-to-neuron signaling in HD patients^93,100–103^, and up-regulated by NA. Nervous system development genes cluster to two groups, up- or down-regulated in vehicle-treated R6/2 striata, perhaps reflective of the complex interplay of genes coordinating neurogenesis, axonogenesis, and glial differentiation, with NA rescuing both groups. Overall, these data support that NA mitigated expression of hundreds of DEGs within HD-related pathways.

The 358 DEGs uniquely elicited by NA cluster within GO pathways important for response to chemicals/small molecules and within HD-related pathways (Extended Fig 4C-D, Extended Table 3). Some of the unique NA-associated gene expression changes could be responses to NA’s binding to CAG slip-outs, responses from targeting repeat instability, or other responses to NA. GO terms within the 47 genes uniquely up-regulated by NA included: Ion binding, Small molecule binding, Cellular response to cAMP, and Negative regulation of fatty acid oxidation (Extended Fig 4B, Extended Table 3). GO terms within the 311 genes uniquely down-regulated by NA included Protein binding, Gated channel activity, Small molecule binding, Synaptic signalling, and Nervous system development (Extended Fig 4C, Extended Table 3).

Overall, these data support that interventional suppression of CAG expansions partially rescued HD-related transcriptionopathy.

### NA does not alter expression of CAG-containing gene transcripts

Although NA is designed to bind to slipped-CAG DNAs, a theoretical possibility is that NA can bind to and alter the expression of CAG-containing gene transcripts. We assessed expression levels of all genes containing at least 3 consecutive CAG units (the shortest stretch that might form a CAG hairpin and be bound by NA), encompassing 385 genes^49^. No significant enrichment of CAG-containing transcripts is evident between NA- and vehicle-treated R6/2 striata (Extended Fig S4D), suggesting that NA did not elicit overt off-target dysregulation of CAG-containing transcripts. This is consistent with our previous finding that NA does not affect transcription across the WT or CAG-expanded *HTT* levels of WT or expanded *HTT* and *ATN1* in HD and DRPLA mice, respectively, or other CAG-containing genes^49,50^.

### NA rescued neuroinflammation pathways which promote striatal cell death

Neuroinflammation pathways can induce neurodegeneration in HD and other brain diseases^109–111^. We assessed levels of Nlrp3, a mediator of the inflammasome, Iba1, a marker for activated microglia (which trigger inflammation), and Caspase-1, a protease which cleaves inflammation and pyroptosis inducing precursors into active peptides thereby regulating inflammation induced cell death. Each of these markers are up-regulated in brains of HD patients and mice^66,99^.

Striatal Nlrp3 signal was significantly elevated in vehicle-treated R6/2 mice relative to WT mice (Fig 5, p < 0.0001, Extended Fig S5A-B, representative whole brain scans and 40x images, Extended Fig S5C, individual mouse quantifications). NA-treated R6/2 striata have significantly less Nlrp3 signal intensity relative to vehicle-treated R6/2 striata (Fig 5, ∼1/2 average reduction, p < 0.0001, Extended Fig S5A-B), although also being significantly higher than WT striata (Fig 5, p = 0.02, Extended Fig S5A-B), suggesting partial rescue.

**Figure 5.**
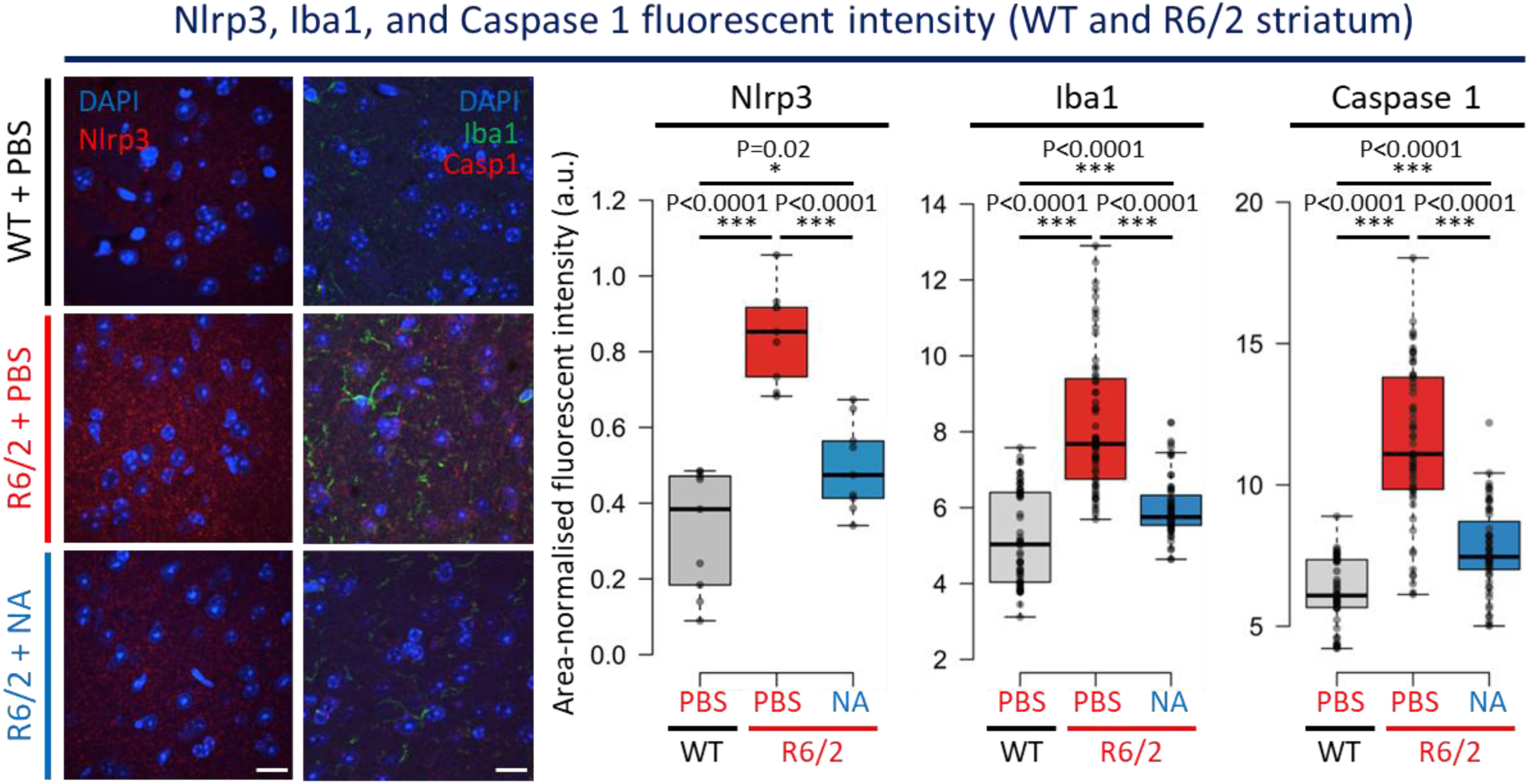
NA diminished neuroinflammation and pyroptosis related pathways in R6/2 striata. Left panels: representative 40x confocal images outlining Nlrp3, Iba1, and Caspase 1 staining in the striatum of 10-week-old vehicle-treated WT mice and R6/2 mice +/−NA. For the leftmost images, red signal denotes Nlrp3 signal and blue signal denotes DAPI-stained nuclei. For the right images, red signal denotes Caspase 1 signal, green signal denoted Iba1 signal, and blue signal denotes DAPI-stained nuclei. Scale bar = 16 μm. Right panels: Nlrp3, Iba1, and Caspase 1 quantifications of area-normalised fluorescent intensity (a.u.) either per whole striatal intensity (Nlrp3) or 40x striatal confocal images of 10-week-old R6/2 mice +/−NA (Nlrp3: n = 3 whole brain slides per mouse and n = 3 mice per condition; Iba1 and Caspase 1: n = 10 images per slide with 3 technical replicates per mouse per condition). Box denotes 1^st^ to 3^rd^ quartiles, black lines denote median, Tukey whisker extents. Statistics: unpaired t-test, *** denotes p<0.0001, * denotes p<0.05.

Nlrp3 signal is also elevated in the frontal cortex (p < 0.0001) and hippocampus (p < 0.0001) of vehicle-treated R6/2 mice relative to WT mice (Extended Fig S5A-B, representative images, and S5C, quantifications). NA significantly reduces fluorescent Nlrp3 signal in both the frontal cortex (p = 0.02) and the hippocampus (p = 0.0002) relative to vehicle-treated R6/2 mice (Extended Fig S5A-C). In the frontal cortex, Nlrp3 intensity in NA-treated R6/2 striata is insignificantly different from WT suggesting complete rescue. In the hippocampus, Nlrp3 intensity was minimally significantly different from WT mice (p = 0.05), consistent with partial rescue.

Significantly higher levels of Iba1 (p < 0.0001) and Caspase-1 (p < 0.0001) are evident in the striatum of vehicle-treated R6/2 mice relative to WT striata (Fig 5, Extended Fig S5D-F for individual mouse quantifications). Consistent with a drop in Nlrp3 signal, Iba1 and Caspase-1 fluorescent intensity was also partially rescued by NA, with significantly less signal intensity relative to vehicle-treated R6/2 striata (p < 0.0001 for both) despite significantly more signal intensity relative to WT mice (p < 0.0001) (Extended Fig S5D-F for individual mouse quantifications).

Overall, these data support that NA reduces neuroinflammation most prominently within the striatum, the brain region most vulnerable to degeneration in HD. NA-mediated reductions in Casp-1 support that NA can impact pathways which promote cell death.

### NA prevented striatal cell depletion and loss of MSN identity markers

Considering the above-noted beneficial effects of NA upon pathways that promote cell death, we assessed if striatal cell loss might also be abrogated. R6/2 mice have been reported to show limited, yet progressive striatal cell loss of 12-25% over 3-12 weeks, where neuron death begins at ∼60 days^53,112^.

We counted up to 1500-2500 striatal cells per mouse per condition within 40X images throughout the striatum. Striata of vehicle-treated R6/2 mice contained significantly fewer cells, with ∼14 less nuclei per image relative to WT mice (Fig 6A, p < 0.0001, Extended Fig S6A, individual mice), a 26% reduction which is consistent with previous reports of neurodegeneration in R6/2 mice^53,112^. NA-treated R6/2 striata showed significantly greater numbers of nuclei than vehicle-treated R6/2 striata, having numbers that were insignificantly different from WT striata (Fig 6A, p < 0.0001, Extended Fig 6A, individual mice). These data support that NA prevents striatal neurodegeneration in R6/2 mice.

**Figure 6.**
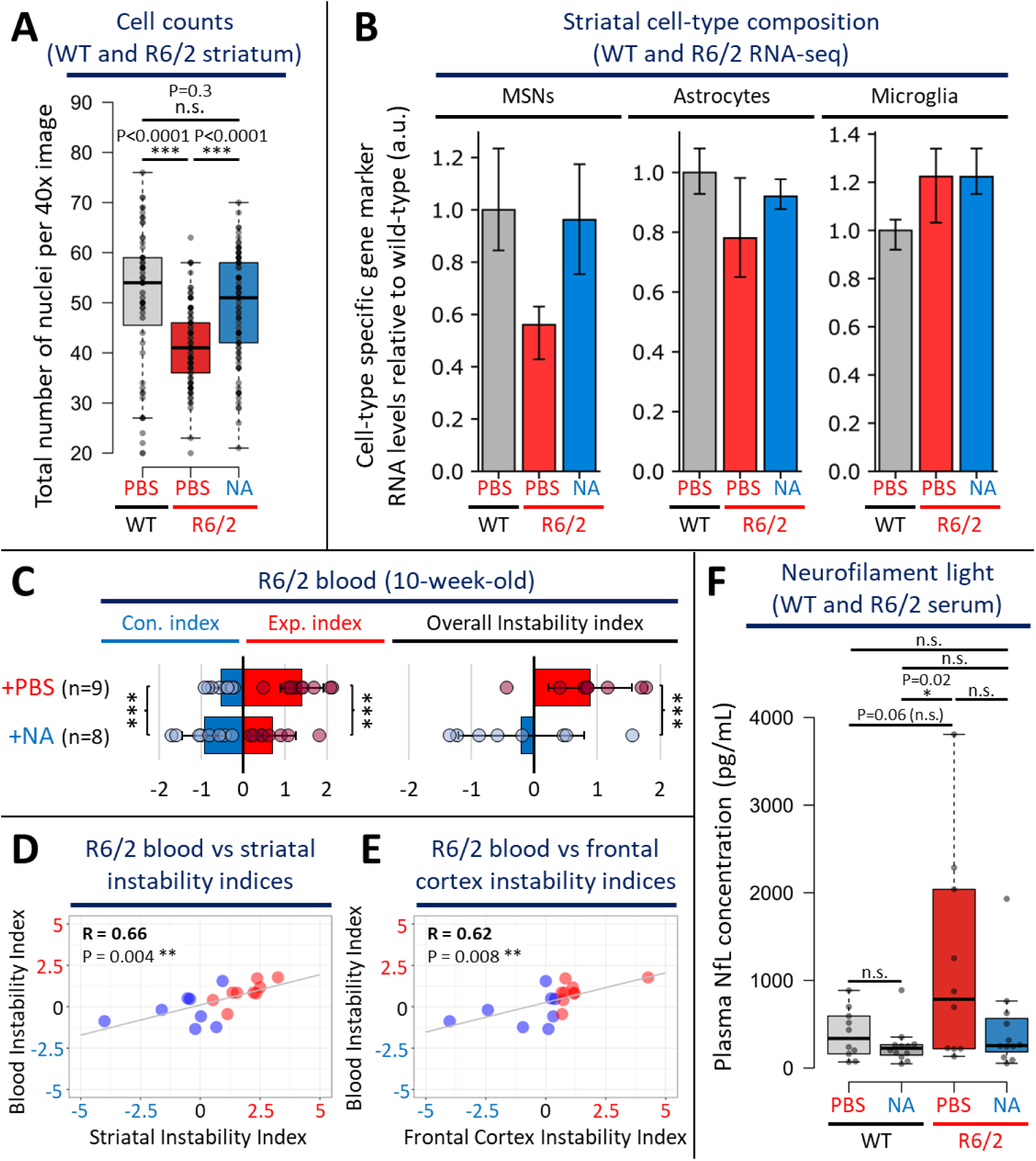
NA prevented striatal cell depletion and loss of MSN identity markers. **A)** quantification of the number of striatal cells in 10-week-old vehicle-treated WT mice and R6/2 mice +/−NA (n = ∼3000-4500 total cells counted over 20-30 40x images per mouse per condition, 2-3 technical replicates per mouse, and 3 mice per condition). Box denotes 1^st^ to 3^rd^ quartiles, black lines denote median, Tukey whisker extents. Statistics: unpaired t-test, *** denotes p<0.0001. **B)** Striatal cell type composition quantifications of n = 7 medium spiny neuron (MSN) and n = 20 astrocyte and microglia cell-type specific markers expression levels derived from 10-week-old vehicle-treated WT and R6/2 HD mice +/−NA striata (n = 3 mice per condition). **C)** Blood repeat expansion indices denoting the average expansion/contraction indices (left panel) and overall instability indices (right panel) in 10-week-old R6/2 mice treated with vehicle (n = 9) or NA (n = 8). Bars denote average instability indices, error bars represent standard deviation from the mean, and circles represent values for individual mice. Statistics: one-tailed Mann-Whitney U test, *** denotes p<0.0001. **D-E)** Pearson correlations comparing blood instability indices to D) striatal and E) frontal cortex instability indices, ** denotes p<0.01. F) quantification of blood serum NfL levels in 10-week-old WT mice treated with vehicle (n = 10) or NA (n = 12) and R6/2 mice treated with vehicle (n = 10) or NA (n = 12). Box denotes 1^st^ to 3^rd^ quartiles, black lines denote median, Tukey whisker extents. Statistics: unpaired t-test, * denotes p<0.05.

As an orthogonal unbiased approach to assess for cell loss, cell-type specific composition was inferred from our striatal RNA-seq data by assessing the relative abundance of cell-type specific RNA transcript levels, as previously demonstrated^113^. Published cell-type-specific RNA-seq data from WT mice of the same background as R6/2 (C57BL/6J) were used to select up to 20 cell-type specific RNA transcript markers per cell-type^90^. Expression levels of individual cell-type-specific markers are reported in Extended Fig 6B.

Vehicle-treated R6/2 striata exhibited ∼1/2 the expression level of 7 MSN-specific marker genes relative to WT striata (Fig 6B, leftmost panel; p < 0.05, 95% confidence interval (CI) overlap). This suggests a depletion or maldevelopment of MSNs in the R6/2 striata. NA-treated R6/2 striata expressed MSN-markers at similar levels to WT, suggesting that NA prevented MSN depletion (Fig 6B). As a confirmatory test, we assessed an additional 19 MSN-specific markers, which again revealed depletion of MSNs in the vehicle-treated R6/2 striata and NA preventing this depletion (Extended Fig 6C-D). The diminished depletion of MSNs by induced CAG contractions is consistent with the reported CAG length dependence of MSN depletion^65,72,73,89^.

Astrocytes specific marker levels (20 astrocyte identity genes), showed mild reductions in vehicle-treated R6/2 striata relative to WT, while NA-treated R6/2 striata exhibit no depletion of astrocytes (Fig 6). This suggests that NA diminished astrocyte degeneration. Microglia specific markers were up-regulated in vehicle-treated R6/2 striata, consistent with increased microglia in HD mice and patients^99,114,115^. NA-treated R6/2 striata showed no effect on microglial specific markers relative to vehicle-treated R6/2 striata. Although NA had beneficial effects on Iba1 protein levels, a marker of activated microglia (Fig 5), the lack of an effect of NA on microglia marker levels suggests that NA may prevent HD-related microglia activation but may not affect microglia accumulation. Despite clear trends, astrocyte and microglia composition was not statistically significant by 95% CI overlap between the 3 groups.

### Blood somatic CAG instability and NfL reflect NAs beneficial effects on brain instability and neurodegeneration

To assess if the beneficial effects of NA could be detected within a readily accessible tissue collectable from living mice, we assessed blood from ICV-delivered NA- or vehicle-treated R6/2 mice. We specifically assessed blood DNA for levels of somatic instability and blood serum for levels of NfL, testing if they could act as surrogate markers of brain CAG instability and neural damage, respectively. Both CAG expansions in blood and NfL in serum have been observed over decades in HD patients but have not been correlated with active expansions in the brain^47^.

Relative to vehicle-treated R6/2 mice (n = 9; instability index: +0.9), blood from NA-treated R6/2 mice (n = 8) demonstrated significant contractions and inhibition of expansions (Fig 6C, instability index: −0.2, p = 0.011, Extended Figure 6F, representative scans). Importantly, blood instability indices significantly positively correlated with instability indices in the striatum (R = +0.66, p = 0.004), frontal cortex (R = +0.62, p = 0.008), and cerebellum (R = +0.53, p = 0.03) of the same mice (Fig 6D-E, Extended Fig 6E). These data support that levels of blood instability reflect levels of brain instability, suggesting that blood instability can act as a surrogate marker for brain instability.

The effects of NA on CAG instability in blood, despite NA being delivered to the CSF via ICV, suggests that while NA is moving into multiple brain regions (indicated by the induced contractions), some portion of NA effluxes from the CSF into general circulation. To assess if ICV-delivered NA can elicit effects in other peripheral tissues, we quantified levels of CAG instability within the gastrocnemius muscle, heart, and liver of NA- and vehicle-treated R6/2 mice (Extended Fig 6F, representative scans). ICV-delivered NA significantly inhibited expansions in the gastrocnemius relative to ICV-vehicle-treated R6/2 mice (Extended Fig 6G). ICV-delivered NA had mild, but insignificant, effects on instability in the heart and liver of R6/2 mice relative to vehicle-treated R6/2 mice (Extended Fig 6G).

To monitor for NA-mediated mitigation of R6/2 neuronal damage/degeneration we quantified levels of blood serum NfL. NfL from blood of R6/2 mice has been observed to increase over 4- to 12-week and older mice, where blood levels reflect those in the CSF and correlated with motor phenotypes^69,70^. Both vehicle- and NA-treated WT and R6/2 mice displayed similar serum NfL levels that were not significantly different, suggesting that NA does not induce neuronal damage in healthy control mice or HD mice (Fig 6C). Vehicle-treated R6/2 mice exhibited significantly higher levels of serum NfL relative to NA-treated WT mice, (Fig 6C, p = 0.02), suggesting that R6/2 mice exhibit higher levels of neuronal damage. Despite having more than ∼2-3-fold higher NfL levels relative to WT mice on average, vehicle-treated R6/2 mice exhibited a wide range of NfL levels; half of the vehicle-treated R6/2 mice having NfL levels that fell within WT ranges while others exhibit ∼5-10-fold higher NfL levels relative to WT (Fig 6C). Due to this intra-group variability, vehicle-treated R6/2 mice were borderline insignificantly elevated relative to vehicle-treated WT mice (Fig 6C, p = 0.06). This is consistent with previous unexplainable variations in NfL levels were reported in R6/2 mice^69^. In contrast, NA-treated R6/2 mice had low and not broadly distributed serum NfL levels, with the mean being insignificantly different from both the broad range of vehicle-treated R6/2 mice and WT mice, suggesting a partial rescue of NfL levels by NA (Fig 6C). Overall, in the context of the cell count and cell composition analyses (Fig 6A & 6B), these NfL data further support that NA prevents neuronal damage/degeneration in R6/2 mice. Furthermore, they suggest that NfL in blood can be considered as a surrogate marker to monitor for changes in neuronal damage while assessing agents targeting somatic instability.

## Discussion

### Insight to preclinical questions

Our preclinical evidence of benefit following interventional *in vivo* modulation of somatic CAG expansions provide validation of instability as a disease-modifying therapeutic target. With multiple anti-expansion approaches in development, the promise of a future interventional treatment is nigh. Our findings begin to lay a framework for designing future anti-somatic instability clinical trials by providing insight to five questions: **1)** Can interventional modulation of somatic expansions effectively lead to benefit of HD-relevant phenotypes in an *in vivo* model? **2)** What *phenotypes* might be affected? **3)** What is the *therapeutic window* to administer treatment? **4)** What is the *time to clinical benefit*? And, **5)** How might we monitor efficacy in readily accessible tissues?

### Interventional modification of somatic expansion modifies HD-related phenotypes

Our findings provide strong evidence for a causal link of somatic expansions to numerous HD-related phenotypes. Expanding on our published proof-of-principle studies^49–51^, here we reveal induced CAG contractions beneficially affect HD-related phenotypic landmarks in one of the most aggressive HD mice, within a treatment window of 5-6 weeks, including: **i)** induced *en masse* brain-wide, muscle, and blood CAG contractions; rescued **ii)** brain-wide toxic polyQ aggregates; **iii)** motor phenotypes; **iv)** muscle strength; **v)** nuclear morphology; **vi)** mRNA nucleocytoplasmic transport; and **vii)** striatal transcriptionopathy; **viii)** aberrant microglial activation; **ix)** aberrant inflammasome; **x)** aberrant pyroptosis; and **xi)** diminished neurodegeneration signatures; **xii)** we also demonstrate that beneficial effects of NA can be monitored through blood, with levels of blood CAG instability reflecting levels of brain instability and decreases in NfL levels reflecting diminished neurodegeneration.

Together these data support that targeting somatic expansions can be clinically beneficial within short time frames, and provide opportunities by which such changes can be monitored in accessible tissues. Thus, our findings provide evidence validating intervention of somatic CAG expansions as a viable avenue for rapid benefit over reasonable timelines^20,116–119^.

### Efficacy and translatability

Our findings provide some insight on the translatability of preclinical animal data to humans. There are key considerations when translating the use of future somatic instability targeting interventions into human expansion-carriers with the expectation of disease-modifying clinical trials. The design of such trials would best rely on evidence-based awareness. The first key factor is efficacy, where an anti-expansion agent would need to slow or suppress active somatic expansions and hopefully induce contractions to a degree such that disease onset and/or progression is beneficially modified in a detectable manner. It is also hoped that such changes would be detectable in the timeframe of a clinical trial.

Here, we used an HD mouse with 120 CAG repeats in all cells at birth, that incur aggressive somatic expansions and disease. 120 is a length inherited by JHD patients (>55 to 350 repeats)^6–11,25^, who incur aggressive somatic expansions and disease^20–22,24–26^. One JHD individual suspected to have inherited (CAG)100 presented with symptoms at 4-years and passed at 6-years and had massive somatic expansions very similar to that arising in (CAG)120 R6/2 mice^26,52,120,121^. 120 is also a length generated by somatic expansions in a large subset of MSNs in brains of humans that inherited (CAG)≥40-55 adult onset HD — where (CAG)80-120 is on the brink of further aggressive somatic expansions upwards of (CAG)1000, leading to extreme pathogenesis and decline in humans^28,29,31^. Delivered to mice for only 6-weeks, NA induced *en masse* CAG contractions of 6-8 repeats and also blocked spontaneous expansions of 10-20 repeats evident in brains of vehicle-treated mice. When considering both contractions and blocked expansions, the net loss induced by NA could be as great as 16-28 repeats, losing as many as 4.6 repeats/week. Such magnitudes of contractions elicited in brains of HD patients, most inheriting (CAG)≥40-55, would bring them closer to or below the reduced penetrance disease threshold of (CAG)≥36-39.

Might such post-zygotic repeat contractions lead to clinical benefits in humans? Contractions of inherited *HTT* expansions of magnitudes similar to those induced herein, have been observed in humans, eliciting rare but insightful cases of diminished disease presentation. For example, while father-to-offspring transmissions are expansion-biased, maternal transmissions are contraction-biased where adult-onset expanded alleles of (CAG)≥40-55 incur losses as large as - 14 repeats down to reduced penetrance or normal lengths^122–128^. Moreover, the human contraction-bias transmitted by mothers is stronger in daughters than sons^122,123^, which, based upon the absence of a parent age-effect, and upon HD mouse and human mechanistic data support post-zygotic contraction events^129–133^. Thus, HD CAG contractions can naturally arise yielding clinical benefits.

Clear human evidence exists of post-zygotic contractions having clinical benefit. In one illuminating example, Nørremølle *et al* reported monozygotic male twins discordant for HD presentation, where the unaffected twin had incurred a post-zygotic mosaic contraction of (CAG)47 down to (CAG)37, while the HD-affected twin inherited (CAG)47^134^. The −10 repeat contraction was prominent in hair root cells which, like the brain, arise from the ectoderm, suggesting the HD-free twin would have the reduced penetrance allele (CAG)37 rather than the fully-penetrant (CAG)47 allele in vulnerable brain cells^135^. In this *“experiment of nature”*, somatic contractions of magnitudes similar to those induced by NA, even at mosaic levels, provided at least 14 years of disease-free living. The affected twin showed motor, cognitive and behavioural symptoms at 30 years of age, at 32 was dismissed from employment they had held for 15 years, and died at 44 from severe HD in 2013. In contrast the CAG-contracted twin was still HD-free over this same time period and is currently alive with unknown clinical state some 26 years after their affected twin began to decline (*personal communication*, Anne Nørremølle, Jørgen E. Nielsen, and Lena Elisabeth Hjermind)^134^. Other clinically discordant HD twins have been reported which may be due to variant levels of somatic instability^136–142^. Clinical benefits in humans incurring CAG contractions of small magnitudes, even if to mosaic levels, further supports targeting somatic expansions as a disease-modifying therapy.

### Intervention in tissues actively incurring somatic expansions

Our findings support the expectation that the impact of an anti-somatic expansion agent would likely be most effective in brain regions incurring the fastest expansion rates. Anti-expansion agents will target entities involved in actively occurring expansions, including MSH3, PMS1, FAN1 and DNA expansion intermediates like slipped-DNAs. The mechanism of action by which NA induces contractions depends upon the same factors required for natural somatic expansions (slipped-DNAs, MSH3, FAN1, and transcription across the expanded repeats) ^49–51^— which further strengthens NA’s specificity for the actively unstable mutant repeat and for cells incurring the fastest somatic expansions. Supporting this, NA-induced contractions more in the faster expanding striatum, followed by the intermediate expansions in the frontal cortex, and least effective in the relatively stable cerebellum. This fuels the expectation that future anti-somatic expansion agents aimed at these factors (MSH3, PMS1, FAN1, and *mHTT* transcription^44,45,143–146^), will likely be more effective in cells incurring the fastest expansions.

### Repeat lengths, intervention timing, and windows of impact

Our findings, in conjunction with other findings, shed light on a repeat length range susceptible to interventional repeat modulation. Repeat length thresholds of extreme somatic expansions and extreme transcriptionopathy in human HD postmortem brains are (CAG)∼80-100 and (CAG)>150, respectively^31^. Similar thresholds are suggested in HD mice. For example, genetic ablation of *Msh3* blocks expansions in both *Hdh*^Q111^ mice with (CAG)100 and zQ175 mice with (CAG)185-190, yet succeeds to delay polyQ aggregate levels in the (CAG)100 but not (CAG)175 mice^147–149^. Similarly, genetically ablating *Msh3* or *Pms1* in Q140 (CAG)140 mice, isogenic to zQ175 (CAG)185, blocks CAG expansions and successfully rescues (or prevents) aggregates, and other deficits^46^. Together, these studies suggest a threshold between (CAG)140-185 for frank disease manifestation in HD mouse models^148^. These animal studies, based on pre-zygotic genetic ablation, coupled with our interventionally NA-induced contractions in R6/2 (CAG)120 mice, suggest that in a clinical setting targeting somatic expansions could be beneficial but should be initiated as early in life as possible to prevent early neuronal death or dysfunction of cells whose expansions have surpassed the suggested pathological threshold of (CAG)150^148,149^.

Our findings might suggest that early premanifest administration of a possible anti-somatic expansion agent could be an impact window allowing possible detectable benefits. Here, NA was delivered to young (4-week-old) R6/2 (CAG)120 mice for 6-weeks. This delivery window coincided with the onset and rapidly increasing rate of somatic CAG expansions, also coincident with the onset and rapid progression of HD-related phenotypes^52,120^. NA’s ability to elicit robust improvements of multiple HD-related phenotypes over this very short time course, supports the possibility of speedy benefits by interventionally targeting somatic CAG expansions. We note that genetic ablation of *Msh3* or *Pms1* in full-length HTT mice (*Hdh*^Q111^, Q140, zQ175) had times to benefit spanning 6-20 months, and may not be detectable with haploinsufficiency^46,147–149^. This may in part be due to the less aggressive aggregation, transcriptionopathy, and phenotypes in full-length HTT mice relative to the rapidly progressing attributes in N-terminal HD mice^55,61,65,89,91,92^.

As noted above, active somatic expansions must be occurring for effective target engagement. The earliest time when somatic expansions begin in humans is not known. They are not present in human HD fetal tissues, but they are evident in brains of premanifest HTT-expansion carriers as young as 44-years of age^31^. While tracts with >100 CAG units were present due to active expansions in striatum of premanifest carriers, they were fewer than those in manifest HD patient brains — again supporting early intervention to prevent rapid accumulation of neurons with larger somatic expansions of (CAG)100-800+, dramatic shifts of dysregulated transcriptomes, and patient decline^31^. Ongoing CAG expansions in blood of premanifest *HTT*-expansion carriers decades before motor onset reveals the tolerability of active somatic expansions up to a limit^31,47^. While fetal brains of human and mouse expansion carriers are devoid of somatic expansions^20,150^, the stable inherited expansion at this prenatal age is not without detrimental neurodevelopmental effects^73,151–153^. As such, the safe modulation of expansions will require balancing of the neurodevelopmental and expected neurodegeneration aspects such that the optimal time for delivery of a possible anti-expansion therapy may vary between individuals^20,116,118,119,154,155^. Natural history studies of *HTT*-expansion carriers also support early administration of possible disease-modifying treatments to premanifest individuals. Our findings, coupled with other mouse interventional studies^156,157^, further support early and/or a longer administration periods and/or more impactful effects upon somatic instability may be optimal for therapeutic benefits.

### Modes of intervention

Our findings, relative to other interventional studies, shed light on the efficacy of the mode of intervention. The efficacy of a possible anti-somatic expansion agent will depend upon the length of the repeat, whether it is actively incurring expansions, and whether the agent slows or ablates expansions, or induces contractions.

Genetic ablation of DNA repair proteins can slow or arrest somatic expansions. Current targets to modulate somatic CAG expansions for therapeutic benefit include MSH3, FAN1, and PMS1, known to act in the formation and processing of slipped-CAG DNAs^28^. HD mice born genetically ablated for MSH3 or PMS1 have ablated or slowed somatic expansions relative to repair-proficient HD mice^46,147,148^. HD mice genetically ablated for FAN1 show increased hyper somatic CAG expansions^158^. To this degree, reducing the activity of MSH3 or PMS1, or increasing the activity of FAN1 would, at maximal efficacy, be expected to ablate somatic expansions. Until recently, the only published benefit of modulating somatic expansions was limited to diminishing polyQ aggregates^49,147^. Q140 (CAG)140 mice genetically null for *Msh3* or *Pms1* showed ablation or slowing of somatic striatal CAG expansions, with blocked or slowed polyQ aggregate formation, respectively^46^. The *Msh3*-null mice also showed reduced striatal synaptic marker loss, astrogliosis, transcriptionopathy, and open field test deficits^46^. These benefits were evident only in mice homozygously-but not heterozygously-deficient for *Msh3* or *Pms1*, and took 6-20 months for the benefits to be detectable^46^. These findings support benefits from inherited genetic deficiencies of MSH3 or PMS1.

Post-natal *in vivo* intervention of somatic expansions is limited to only a few studies, with the exception of two studies^49,50^, all induced slowing or speeding of somatic expansions but not stopping or reversing them^45,156,157^. Adeno-associated virus mediated CRISPR–Cas9 genome editing (likely bi-allelic) of *Msh3* in striata of *Htt*^Q111^ (CAG)112-119 mice after 6-months showed dramatic slowing of somatic expansions^45^. Varying degrees of slowing or speeding somatic expansions can arise depending upon which DNA repair protein is targeted^45^. Surprisingly, CRISPR–Cas9 targeting of 63 DNA repair proteins or genetically ablating nine GWAs-identified HD age-of-onset modifiers, revealed many that slowed, sped, or had no effect upon somatic expansions, but none induced contractions^45,46^.

Slowing somatic expansions interventionally may need to be sustained for long periods to detect clinical benefits, as noted above. Knocking-down MSH3 or PMS1 by 60-80% for 7-11-weeks in (CAG)110-125 cell models, slowed expansions to varying degrees^144,145^. Two *in vivo* interventions by ICV delivery of an anti-*Msh3* siRNA to 3-month old *Hdh*^Q111/+^ (CAG)109-111 mice slowed but did not arrest striatal CAG expansion rates over 3-months post-delivery^156,157^. This intervention had no impact upon mHTT aggregation^156^, which contrasts with the delay of aggregates following arrest of somatic expansions in 5-month old *Hdh*^Q111/+^ mice born genetically ablated for *Msh3*^147^. Neither of these interventional or genetic ablation studies reported neuropathological or motor benefit. Extended treatment courses and/or high levels of knockdown/inhibition may be required to detect phenotypic benefits.

Induced CAG contractions, in the face of spontaneous expansions, over brief intervention periods (6-weeks), as reported here, can rapidly diminish deleterious effects of many HD-related phenotypes (see above section *“Interventional modification of somatic expansion modifies HD-related phenotypes”*). As such, benefits of a much-anticipated disease-modifying therapy, with slowing or ablating somatic repeat expansions, or inducing contractions, can occur within reasonable timeframes.

### Assessing impact of interventionally modified somatic expansions in an accessible tissue

Monitoring target engagement and treatment efficacy in an anti-somatic expansion disease-modifying clinical trial is needed. Since blood-derived somatic CAG expansions could be detected decades before disease onset and tracked with early and longitudinal changes monitorable by the HD Integrated Staging System (HD-ISS)^154^, including brain atrophy (NfL), suggests blood DNA expansions may be a surrogate marker for somatic expansions in the brain^47^. This connection of somatic expansions in blood and brain has not been demonstrated in the same organism. Were this suggested link of somatic expansions in blood a proxy to expansions in the brain validated, this would present the blood as a possible source of target engagement for anti-expansion agents that contact the blood. Since the levels of *HTT* CAG tract expansions in blood and skeletal muscle reflect the levels of expansion in multiple brain levels^26^, and that we observed effects of NA upon the levels of CAG instability in blood and muscle, suggests that both accessible tissues may, for some anti-somatic expansion agents, be suitable surrogate markers of target engagement. Our study provides the first evidence supporting the potential utility of blood to monitor the effects of CAG-target engagement.

Our findings show diminished aberrant NfL levels in serum are evident with induced CAG contractions in HD mouse brains and benefits of HD-related phenotypes, consistent with NfL levels being a predictor of neurodegeneration and disease progression in various HD mice^69,70^. That ICV-administered NA led to diminished levels of NfL in blood plasma supports the ability to monitor the expected beneficial effects of a future anti-expansion agent. Our study provides the first evidence supporting the potential utility of blood to monitor the effects of CAG-target engagement.

### Limitations

We recognize the limitations of HD mouse models in that they do not recapitulate either the scale or spectrum of disease phenotypes in HD patients, and that inherited CAG lengths in mice need to be >100 repeats to show HD-related phenotypes, as opposed to the most common inherited repeat range of 40-45 in HD patients. This may reflect either murine-specific tolerability’s or the reduced lifespan available for mice to fully-develop disease.

Similarly, different ageing rates between mice and humans, where life phases and maturational rates of mice do not linearly correlate with humans—being 45- to 150- times faster, depending upon the life phase^159^. This makes it challenging for this, and all mouse preclinical studies, to translate mouse ageing times to human ageing times.

Notwithstanding these limitations, our study demonstrates R6/2 (CAG)120 may be a valuable model to study somatic CAG-expansion driven HD-related phenotypes and how interventional targeting of somatic CAG expansions could modify these disease processes.

### Summary

Our findings provide valuable support favouring efforts to target somatic CAG expansions as a viable disease-modifying therapeutic approach for HD disease and possibly other repeat expansion diseases. These preclinical suggests that a therapeutic window of treatment can begin early in premanifest and early manifest individuals. These proof-of-principle findings provide further support to searches for anti-somatic repeat expansion interventions as disease-modifying therapies for HD and other repeat diseases and suggest that when available these therapies could be effective within time frames of clinical trials.

## Methods

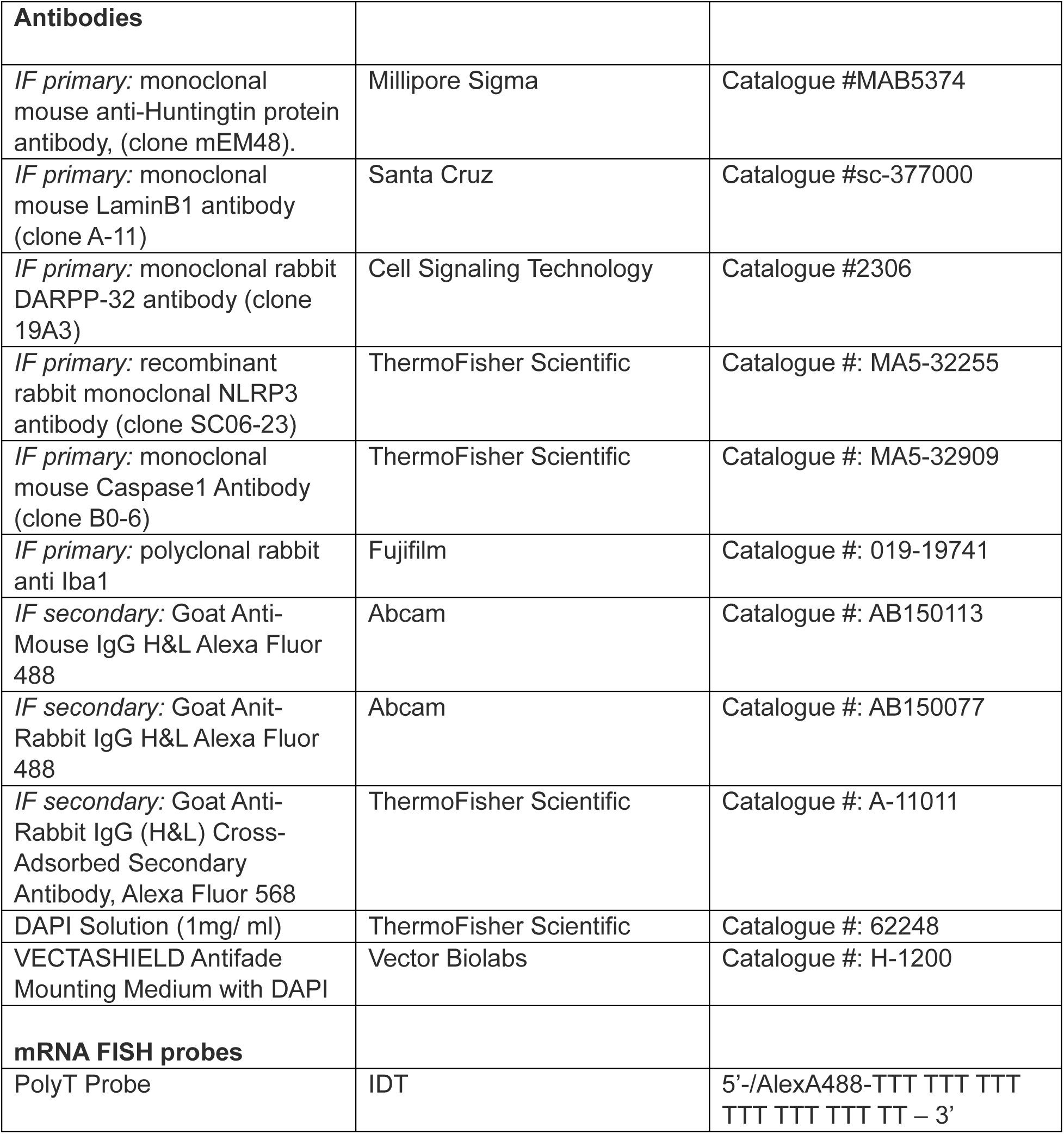

|  |  |  |
| --- | --- | --- |
| **Antibodies** |  |  |
| *IF primary*: monoclonal mouse anti-Huntingtin protein antibody, (clone mEM48). | Millipore Sigma | Catalogue #MAB5374 |
| *IF primary*: monoclonal mouse LaminB1 antibody (clone A-11) | Santa Cruz | Catalogue #sc-377000 |
| *IF primary*: monoclonal rabbit DARPP-32 antibody (clone 19A3) | Cell Signaling Technology | Catalogue #2306 |
| *IF primary*: recombinant rabbit monoclonal NLRP3 antibody (clone SC06-23) | ThermoFisher Scientific | Catalogue #: MA5-32255 |
| *IF primary*: monoclonal mouse Caspase1 Antibody (clone B0-6) | ThermoFisher Scientific | Catalogue #: MA5-32909 |
| *IF primary*: polyclonal rabbit anti Iba1 | Fujifilm | Catalogue #: 019-19741 |
| *IF secondary*: Goat Anti-Mouse IgG H&L Alexa Fluor 488 | Abcam | Catalogue #: AB150113 |
| *IF secondary*: Goat Anti-Rabbit IgG H&L Alexa Fluor 488 | Abcam | Catalogue #: AB150077 |
| *IF secondary*: Goat Anti-Rabbit IgG (H&L) Cross-Adsorbed Secondary Antibody, Alexa Fluor 568 | ThermoFisher Scientific | Catalogue #: A-11011 |
| DAPI Solution (1mg/ ml) | ThermoFisher Scientific | Catalogue #: 62248 |
| VECTASHIELD Antifade Mounting Medium with DAPI | Vector Biolabs | Catalogue #: H-1200 |
| **mRNA FISH probes** |  |  |
| PolyT Probe | IDT | 5'-/AlexA488-TTT TTT TTT TTT TTT TTT TT – 3' |

### Mice

R6/2 transgenic mice and WT littermates were obtained from the Jackson Laboratory (Strain: #006494, [Tg(HDexon1)62Gpb]). R6/2 transgenic mice carry an *HTT* exon-1 sub-fragment containing 120-121 CAG repeats. The mice were purchased form the Jackson Laboratory, shipped to the Hospital for Sick Children Lab Animal Service, and were group-housed in cages, with each cage housing 2-5 mice. The housing conditions were maintained at a constant temperature of 22 °C, with a 12-hour light/dark cycle, and food and water were provided ad libitum. All procedures were approved by the Hospital for Sick Children’s Animal Care and Use Committee and were conducted in accordance with the guidelines of the Canadian Council on Animal Care and the National Institutes of Health.

### ICV pump implantation

Intracerebroventricular (ICV) administration of NA or vehicle (phosphate-buffered saline; PBS) was performed with an ALZET® osmotic pump (model #2004; Alzet Corp, Palo Alto, CA, USA) as per the manufacturer’s instructions. Briefly, 4-week-old mice were anesthetized with isofluorane and the osmotic pump, filled with NA (500 uM stock) or sterile PBS, was implanted in a subcutaneous pocket that was formed on the back of the mouse. The catheter was placed into the left lateral ventricle found using the following coordinates from the bregma: AP: +/−1.0 mm, ML: −0.2 mm, and DV: −2.0 mm (x, y, z). NA or PBS was continuously infused at 0.2 μL/hour for the duration of the experiment.

### Fragment Length Analysis

Genomic DNA was isolated from mouse brain tissues using the Gentra Puregene Kit (Qiagen, Valencia, CA). DNA from 2-week-old mouse tail and ear notches was extracted to determine the inherited CAG length of the mice. For repeat length analysis, genomic DNA was amplified using Platinum SuperFi II PCR Master Mix (ThermoFisher Scientific, catalogue #12368050) with the following primers: Forward primer: FAM-labelled 5’-ATGGCGACCCTGGAAAAGCTGATGAA-3’ and Reverse primer: 5’-GGCGGCTGAGGAAGCTGAGGA-3’. The PCR cycles were as follows: 1) 1X 98°C for 30 seconds 2) 28X (98°C for 10 seconds, then 62°C for 10 seconds, then 72°C for 30 seconds), 3) 72°C for 5 minutes, 4) infinite hold at 4°C. The fluorescently labelled PCR products were then processed by capillary gel electrophoresis with size markers on a SeqStudio Flex Genetic Analyzer (ThermoFisher Scientific). Peak Scanner software Version 3.0 (ThermoFisher Scientific) was used to visualize and size repeat lengths. Instability index calculations were done as previously described^160^.

### Immunofluorescence

Whole mouse brains were fixed in 4% paraformaldehyde for 12-16 hours, embedded in paraffin, and 5μm sagittal sections were obtained. Xylene was used to deparaffinize the sections, which were rehydrated in serial dilutions of ethanol (100 x2, 95, and 70%) followed by ddH2O and 1X PBS for 5 minutes each. Antigen retrieval was performed by heating slides in a steamer in 0.01M citrate buffer (pH 6.0) for 20 mins. Sections were blocked with 10% normal goat serum in 1xPBS + 10% Triton X-100 for 1 hour at room temperature.

Sections were then incubated with primary antibodies in blocking solution overnight at 4C. Slides were washed for 5 minutes 3x with 1xPBS-T, and incubated with secondary antibodies in 1XPBS-T for 1 hour at room temperature, then washed for 5 mins 3x with 1xPBS-T. Nuclei were stained with 0.2 μg/mL DAPI in 1XPBS-T and mounted with VECTASHEILD antifade mounting media. Images were acquired using a IX83-F3 Olympus confocal microscope. Fluorescent signal was analysed and quantified using ImageJ and HALO image analysis platform.

### Mouse behavior

Five weeks after pump implantation and treatment initiation, R6/2 transgenic mice and age-matched wild-type mice were subjected to four different behavioral tests. The order of testing progressed from the least to the most stressful (e.g., open field → hindlimb clasping → wire hanging → rotarod). A 3-day interval was maintained between each behavioral test to minimize any potential carryover effects. All experiments were conducted during the light phase and were performed and analyzed in a blinded manner. The sample sizes were determined based on established standards in the field and previous experience with phenotype comparisons. No statistical methods were used to predetermine the sample size.

### Open field

Open field test was conducted as previously described with minor modifications^161^. Briefly, each mouse (R6/2 + PBS, *n* = 12, R6/2 + NA, *n* = 16, WT + PBS, *n* = 10, and WT + NA, *n* = 7) was positioned at the center of an opaque, white-hued open field arena (46 cm × 46 cm × 20 cm). The locomotive behavior was video-recorded for a duration of 10 minutes, followed by an automated analysis using LimeLight software (Actimetrics, Wilmette, IL). For the analysis of locomotor activity, the total distance traversed in the open field arena was calculated. Two-way ANOVA with Holm-Šídák post-hoc tests were used for statistical analysis. All individual data points are plotted with crossbar denoting mean ± SEM.

### Wire hanging

Wire hanging test was conducted as previously described with minor modifications^162^. Briefly, each mouse (R6/2 + PBS, *n* = 13, R6/2 + NA, *n* = 13, WT + PBS, *n* = 5, and WT + NA, *n* = 7) was allowed to grip and suspend itself from a horizontally positioned wire (2 mm in diameter) using its forelimbs, and the duration of the hang was captured on video until the subject fell. The time spent hanging on the wire was quantified to assess muscle coordination and strength. Two-way ANOVA with Holm-Šídák post-hoc tests were used for statistical analysis. All individual data points are plotted with crossbar denoting mean ± SEM.

### Rotarod

Rotarod task was conducted as previously described with minor modifications^60^. Briefly, each mouse (R6/2 + PBS, *n* = 10, R6/2 + NA, *n* = 12, WT + PBS, *n* = 11, and WT + NA, *n* = 6) was placed onto an accelerating Rotarod system (Med Associates, VT) to assess motor coordination and motor learning. The mouse underwent a 3-day training period, consisting of 4 trials per day on the Rotarod. Each trial had a maximum duration of 5 minutes, and a 10-minute inter-trial interval was provided for recovery and rest. During each trial, the speed of rotation on the Rotarod accelerated linearly from 4 r.p.m. to 40 r.p.m. The latency of mouse retention on the rod before falling was automatically recorded via an infrared motion sensor below the rod. The latency values of the 4 trials conducted each day were averaged and analyzed. Repeated measures two-way ANOVA with Holm-Šídák post-hoc tests were used for statistical analysis. Data plotted as mean ± SEM.

### Hindlimb clasping

Hindlimb clasping was conducted as previously described with minor modifications^163^. Briefly, each mouse (R6/2 + PBS, *n* = 14, R6/2 + NA, *n* = 14, WT + PBS, *n* = 6, and WT + NA, *n* = 7) was suspended by its tail and video-recorded for 30 seconds. Hindlimb clasping scores were calculated to evaluate cerebellar ataxia as follows: Score zero, when both hindlimbs splayed outward and away from the abdomen; Score one, when one hindlimb retracted toward the abdomen; Score two, when both hindlimbs retracted toward the abdomen; Score three, when both hindlimbs fully retracted toward the abdomen and were touching each other; Score four, when forelimbs were touching both fully retracted hindlimbs. The scores were calculated for each 10-second interval and averaged for each mouse. Two-way ANOVA with Holm-Šídák post-hoc tests were used for statistical analysis. All individual data points are plotted with crossbar denoting mean ± SEM.

### Correlation analysis

Somatic repeat instability index, contraction index, and expansion index in R6/2 HD PBS and NA-treated mice were correlated with phenotypic outcomes in R (version 4.3.3) using a Pearson correlation. The ‘ggplot2’ R package was used to visualize all correlations.

### RNA Fluorescent *In Situ* Hybridization (FISH)

OCT-compound-embedded frozen whole mouse brains were sectioned and 10 uM sagittal sections were obtained. Sections were equilibrated at room temperature for 15 minutes and washed in 1XPBS for 5 minutes 3x to remove OCT. Slides were permeabilized in 0.2% Triton X-100 – PBS for 10 minutes at room temperature and blocked with hybridization buffer (50% formamide, 2X saline-sodium citrate (SSC), 50 uM sodium phosphate, 10% dextran sulfate, and 2 mM vanadyl sulfate ribonucleosides) at 37C for 30 minutes. Slides were then hybridized with 200 uM 5’ labeled Alexa488 PolyT oligonucleotide probes in hybridization buffer at 37C overnight. Slides were then put into stringency washes once with 4X SCC and once with 2X SCC for 10 minutes each at room temperature. Following washes, slides were incubated in 0.1% TritonX100 −2X SCC with 0.2 ug/ml DAPI for 15 minutes at room temperature. Slides were washed twice with 2X SCC for 5 minutes each. Autofluorescence was quenched with 0.25% Sudan Black in 70% ethanol for 15 seconds. Slides were then washed in 2X SCC for 5 minutes and mounted with VECTASHEILD antifade mounting media. Images were acquired using a IX83-F3 Olympus confocal microscope. mRNA nuclear retention was quantified using ImageJ by firstly normalising total mRNA fluorescent intensities per 40x image to the total area of the image (overall baseline signal per area), secondly normalising the fluorescent intensity of each individual nuclei by the nuclei’s area (nuclei-specific mRNA signal per nuclear area), and then thirdly normalising individual nuclear intensities to the normalised full image intensity (assessing the relative intensity of individual nuclear signals over the baseline signal), with higher intensity within the nucleus (i.e. mRNA retention) quantified as higher intensity values.

### RNA-seq library preparation and sequencing

RNA sequencing (RNA-seq) libraries were prepared using the SureSelect XT RNA Direct kit (Agilent Technologies) following the manufacturer’s protocol. Sequencing was performed on an Illumina NovaSeq 6000 platform, and reads were aligned to the mouse reference genome (GRCm38) using DRAGEN Bio-IT software version 3.6.3 (Illumina).

### Gene expression analysis

RNA expression in transcripts per million (TPM) for each gene was quantified by pseudoalignment using kallisto v0.43.0 against the mm10 reference genome^164^.

Principal component analysis (PCA) was performed in Python 3.6.5 using scikit-learn v0.20.0^165^. Differential gene expression (DE) was quantified pairwise between WT, R6/2 PBS, and R6/2 NA treatment groups using DESeq2 v1.30.1 directly from pseudoaligned transcript abundances^166^. A gene was called as DE between two conditions if |log_2_(fold change)| > 0.5 and if the false discovery rate (FDR) q < 0.05, corrected using the Benjamini-Hochberg method^167^. Volcano plots were generated using the ‘EnhancedVolcano” R package and histograms were generated using Python 3.6.5.

### Gene ontology (GO) analysis and heatmaps

GO analysis was carried out using the Gene Ontology Consortium resource (https://geneontology.org/) to identify enrichment of dysregulated genes against a background of 26944 protein-coding genes. GO terms were filtered to only include those with an adjusted p-value (q) of q < 0.05. The ‘pheatmap’ R package was used to visualize heatmaps of genes associated with selected GO terms in mice across different treatment conditions.

### Off-target effects of NA on CAG-containing genes

CAG-containing genes were identified in the Mus musculus mm39 genome by identifying any protein-coding genes with at least 3 consecutive (CAG) repeats, the minimum number of repeats required to form a slip-out to which NA can bind. A Fisher’s exact test was carried out to determine whether dysregulated genes were enriched for CAG repeats. This was visualized as a boxplot using the ‘ggplot2’ R package.

### Cell type composition estimation

Cell type composition changes between conditions were estimated from bulk RNA-seq using a marker gene approach. A published dataset of single-cell RNA-seq performed in WT and R6/2 mouse striatum (GEO accession #GSE152058) was identified in the literature and used to algorithmically define optimal marker genes via pseudobulk analysis^90^. Keeping WT and R6/2 conditions separate, transcript counts for each gene were summed across all cells within each annotated cell type, and a pseudocount was added to each gene to reduce the impact of sparse data on marker gene selection. Neuronal cell subtypes were merged into broader categories of MSN (containing iMSN and dMSN) and other neurons (containing GABAergic, cholinergic, and parvalbumin interneurons, Foxp2+ neurons, neural progenitor cells). Ependymal, mural, and endothelial cells were merged into the category “Other”. For each cell type in each condition, TPM values for each gene were calculated from the summed gene counts.

Marker genes were selected by an entropy minimization approach. Genes that were undetected in at least one mouse striatum sample in our bulk RNA-seq study were excluded. For each gene, the TPMs in each pseudobulk cell type from WT mice were converted into a discrete probability distribution (normalized to sum to 1). Entropy (H) was calculated as the following for each gene:

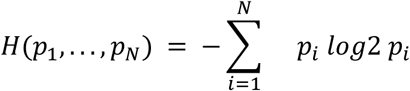

Specificity (s) was defined as the following and also calculated for each gene:

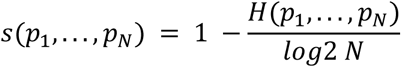

For each cell type, the top 20 most specific genes with minimum expression of 50 TPM in that cell type and >40% minimum specificity were chosen as markers for estimating cell type composition of the bulk RNA-seq samples in this study. For MSNs 7 genes were found to meet these criteria (Tspear, Slit3, Sh3rf2, Dlgap2, Lrtm1, Zfp831, Rbfox3).

To estimate the relative change in cell type composition compared to control mice, the expression in TPM of each gene was normalized by its mean expression in WT mouse samples, and the mean normalized expression of all markers for each cell type was calculated for each RNA-seq library. For MSNs, an additional analysis using the same single-cell RNA-seq reference was performed with an independent set of 19 SPN marker genes defined in the literature to confirm the result^90^.

### NfL quantification

NfL concentrations (pg/mL) in mouse plasma were measured using a commercially available Neuro-4-Plex B (N4PB) kit on the SIMOA HD-X analyser platform following the manufacturer’s instructions (Quanterix, Billerica, MA, catalog #103345). Samples were diluted 1:40 with dilutant provided within the kit or 1:80 if there was insufficient volume. Controls and diluted samples were loaded into the HD-X analyser and analysed in duplicate. The intra-assay coefficient of variance for each sample’s duplicate measurement was < 10%. All sample concentrations were above the lower limit of quantification for the assay (0.500 pg/mL).

## EXTENDED FIGURES

**Extended Figure 1.**
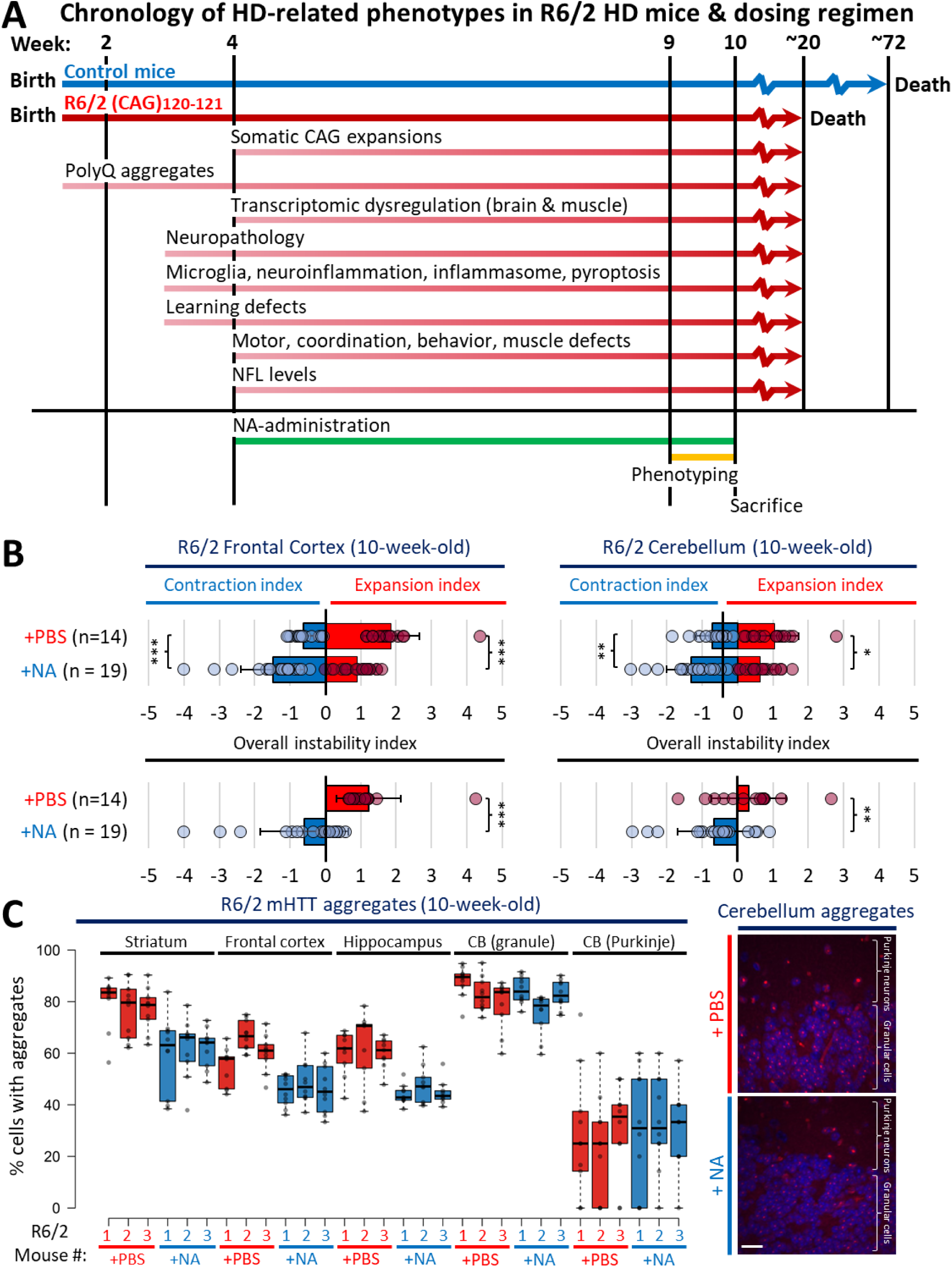
NA modulates somatic instability and aggregates in multiple brain regions. **A)** Detailed summary of treatment regimen, treatment timeline, phenotypic assessments, and tissue harvesting for mice used in this study, in the context of previously published disease presentation chronology in R6/2 mice. B) Frontal Cortex (left panels) and cerebellum (right panels) repeat expansion indices denoting the average expansion/contraction indices (top panels) and overall instability indices (bottom panels) in 10-week-old R6/2 mice treated with vehicle (n = 14) or NA (n = 19). Bars denote average instability indices, error bars represent standard deviation from the mean, and circles represent values for individual mice. Statistics: one-tailed Mann-Whitney U test, *** denotes p<0.0001, ** denotes p<0.01. D) Right panel: representative 40x confocal images outlining mutant poly-glutamine aggregates in the cerebellum of 10-week-old R6/2 mice +/−NA. Red signal denotes mutant HTT (mHTT) poly-glutamine fragment aggregates, blue signal denotes DAPI-stained nuclei. Labels designate Purkinje neurons from granule cells. Scale bar = 16 μm. Left panel: Individual mice per group quantification of percentage of cells with nuclear poly-glutamine aggregates relative to all nuclei per 40x image in the striatum, frontal cortex, hippocampus, and cerebellum (CB) of 10-week-old R6/2 mice +/−NA (n = 10 images per mouse, ∼300-500 cells assessed total per mouse, with n = 3 mice per condition). Box denotes 1^st^ to 3^rd^ quartiles, black lines denote median, Tukey whisker extents.

**Extended Figure 2.**
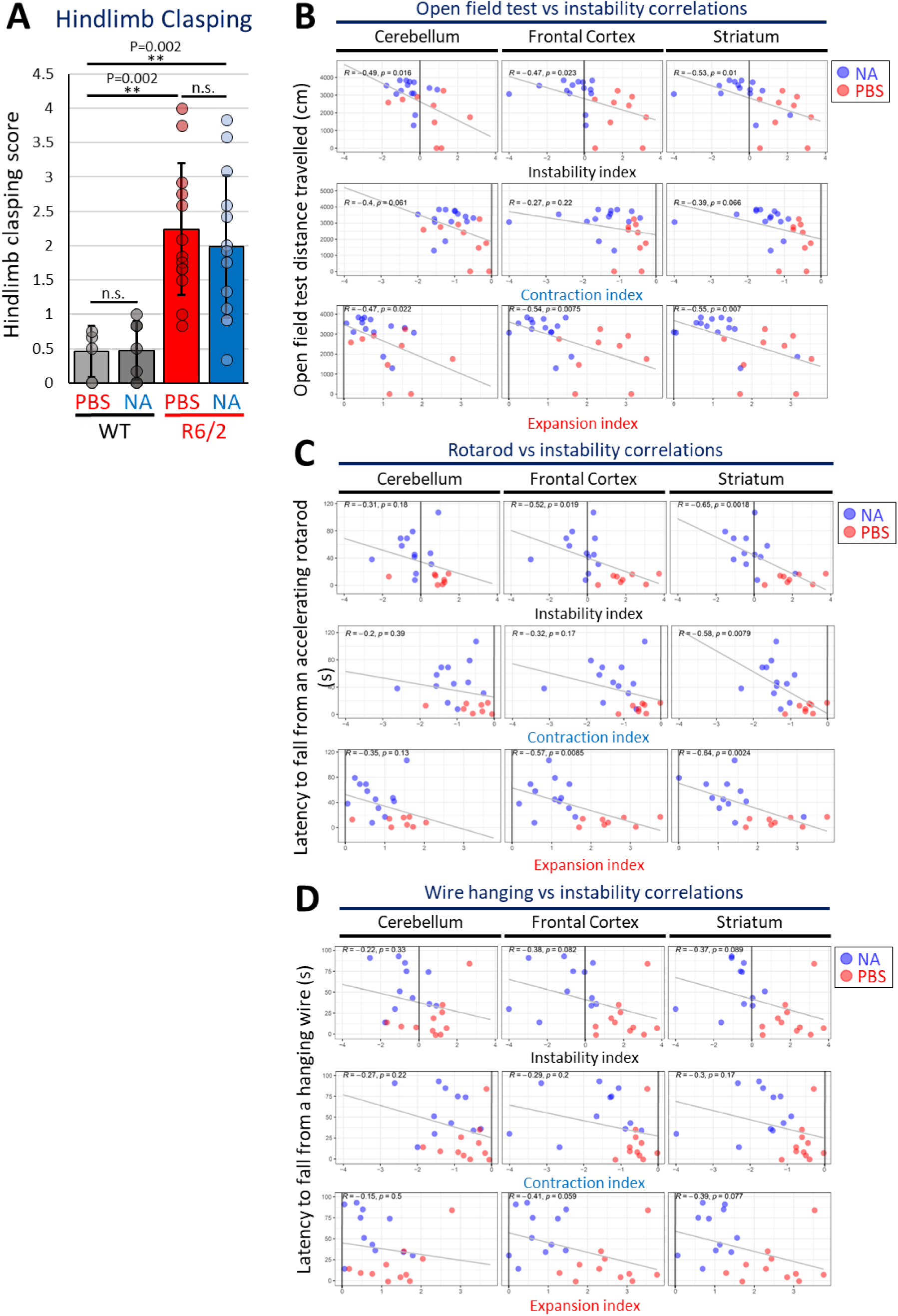
Somatic instability in the striatum and frontal cortex correlates with motor phenotype severity. **A**) R6/2-PBS mice exhibit an increased hindlimb clasping score compared to WT-PBS and WT-NA mice, whereas R6/2-NA mice do not show recovery from the phenotype (two-way ANOVA; Genotype effect: *F*_1,37_ = 20.47, *P* < 0.0001, Treatment effect: *F*_1,37_ = 1.42, *P* = 0.2416; *n* = 6 (WT-PBS), 7 (WT-NA), 14 (R6/2-PBS), 14 (R6/2-NA)). Results are shown as mean ± SEM. ** denotes p<0.01, n.s. is not significant. **B)** Pearson correlations comparing cerebellar, frontal cortex, and striatal instability, expansion, and contraction indices to distance travelled in an open field. **C)** Pearson correlations comparing cerebellar, frontal cortex, and striatal instability, expansion, and contraction indices to latency to fall from an accelerating rotarod. **D)** Pearson correlations comparing cerebellar, frontal cortex, and striatal instability, expansion, and contraction indices to latency to fall from a hanging wire.

**Extended Figure 3.**
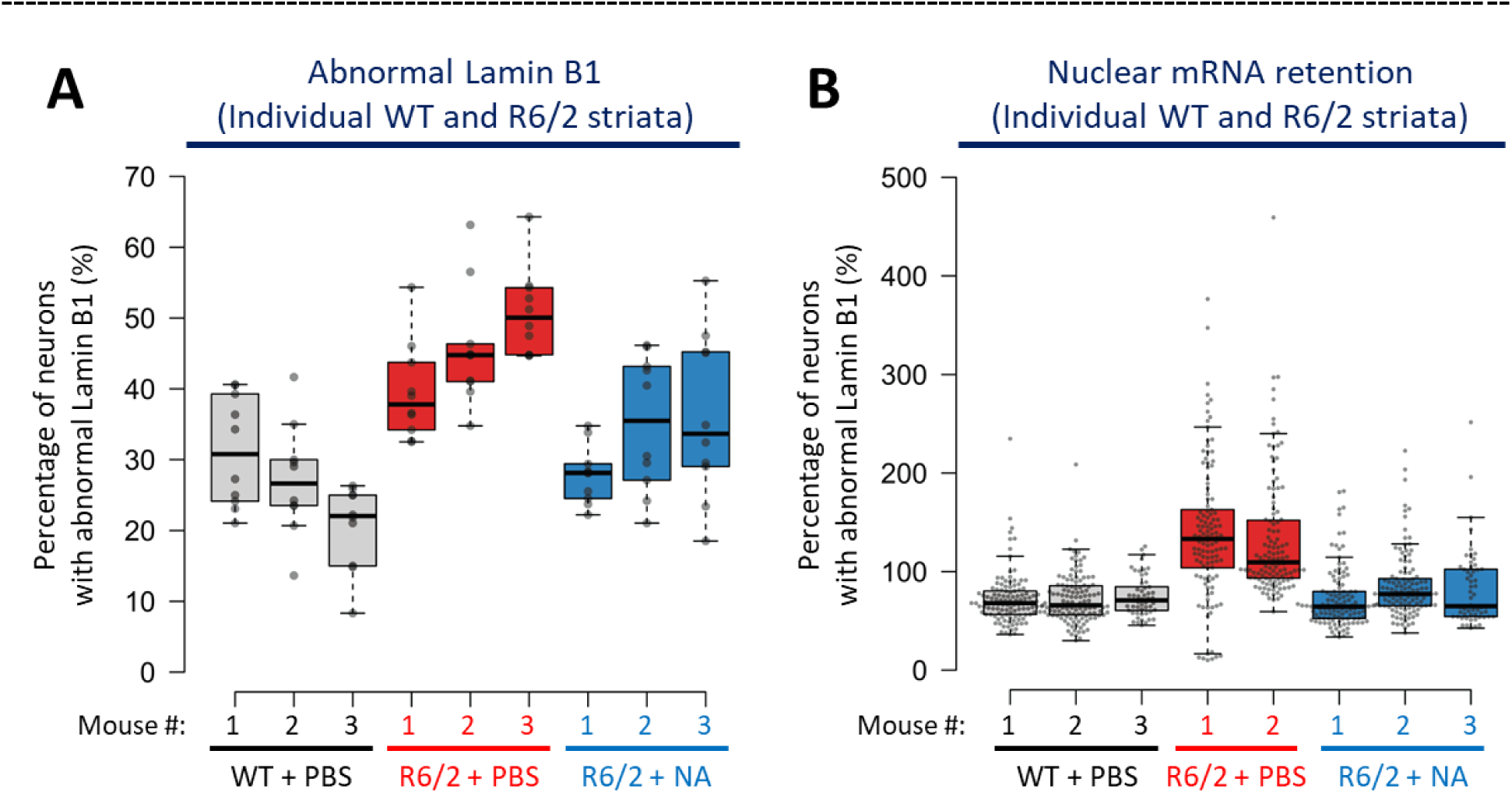
NA improves nuclear envelope morphology and mRNA nuclear export in R6/2 HD striata. **A)** Individual mice quantification of percentage of cells with abnormal Lamin B1 morphology relative to all nuclei per 40x image in the striatum of 10-week-old R6/2 mice +/−NA (n = 10 images per mouse per condition, ∼350-550 cells assessed total per mouse, with n = 3 mice per condition). **B)** Individual mice quantification of normalised nuclear mRNA fluorescence for cells in the striatum of 10-week-old R6/2 mice +/−NA (n = ∼125 individual cells assessed over 2 technical replicates per mouse, each replicate assessing all nuclei within 8-10 40x images, n = 3 WT mice and NA-treated R6/2 mice and n = 2 PBS-treated R6/2 mice). Boxes denotes 1^st^ to 3^rd^ quartiles, black lines denote median, Tukey whisker extents.

**Extended Figure 4.**
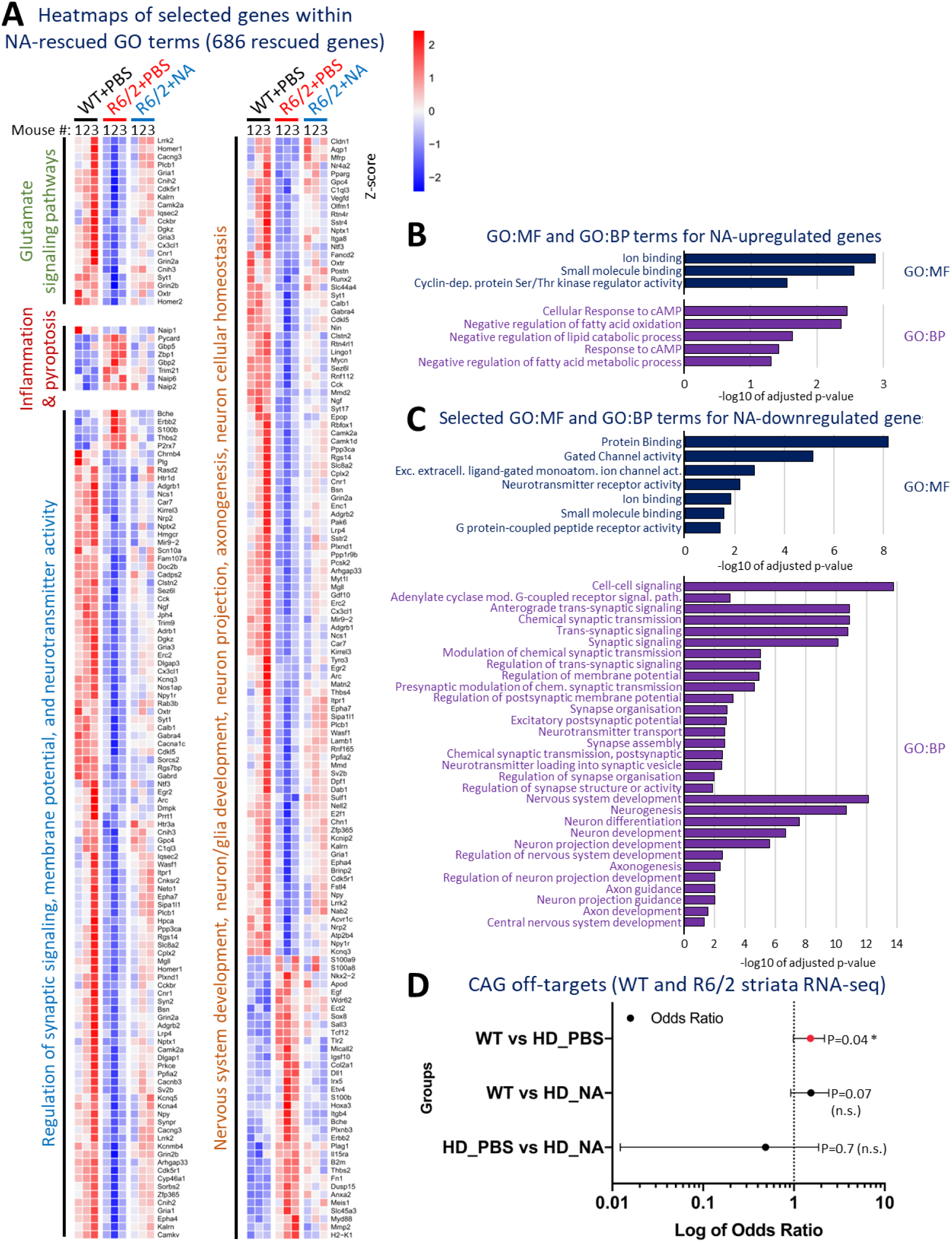
NA mitigates disease-associated transcriptionopathy in R6/2 HD striata. **A)** Heatmaps outlining the expression of all genes that were associated with the 4 major selected HD-relevant GO terms (glutamate signalling, pyroptosis, synaptic signalling, nervous system development). Individual columns represent the expression profile of each mouse per group (n=3 mice per group). Gene expression is represented by Z-scores where a blue represents downregulation relative to the mean expression and red represents upregulation relative to the mean expression. **B)** GO: molecular function (GO:MF) and GO: biological function (GO:BP) terms for the genes uniquely upregulated in NA-treated R6/2 striata vs WT relative to vehicle-treated striata vs WT. **C)** GO: molecular function (GO:MF) and GO: biological function (GO:BP) terms for the genes uniquely downregulated in NA-treated R6/2 striata vs WT relative to vehicle-treated striata vs WT. **D)** Fischer’s test assessing for the enrichment of differentially expressed CAG-repeat containing transcripts in the striata of R6/2 HD mice +/−NA relative to each other and relative to vehicle-treated WT mice striata.

**Extended Figure 5.**
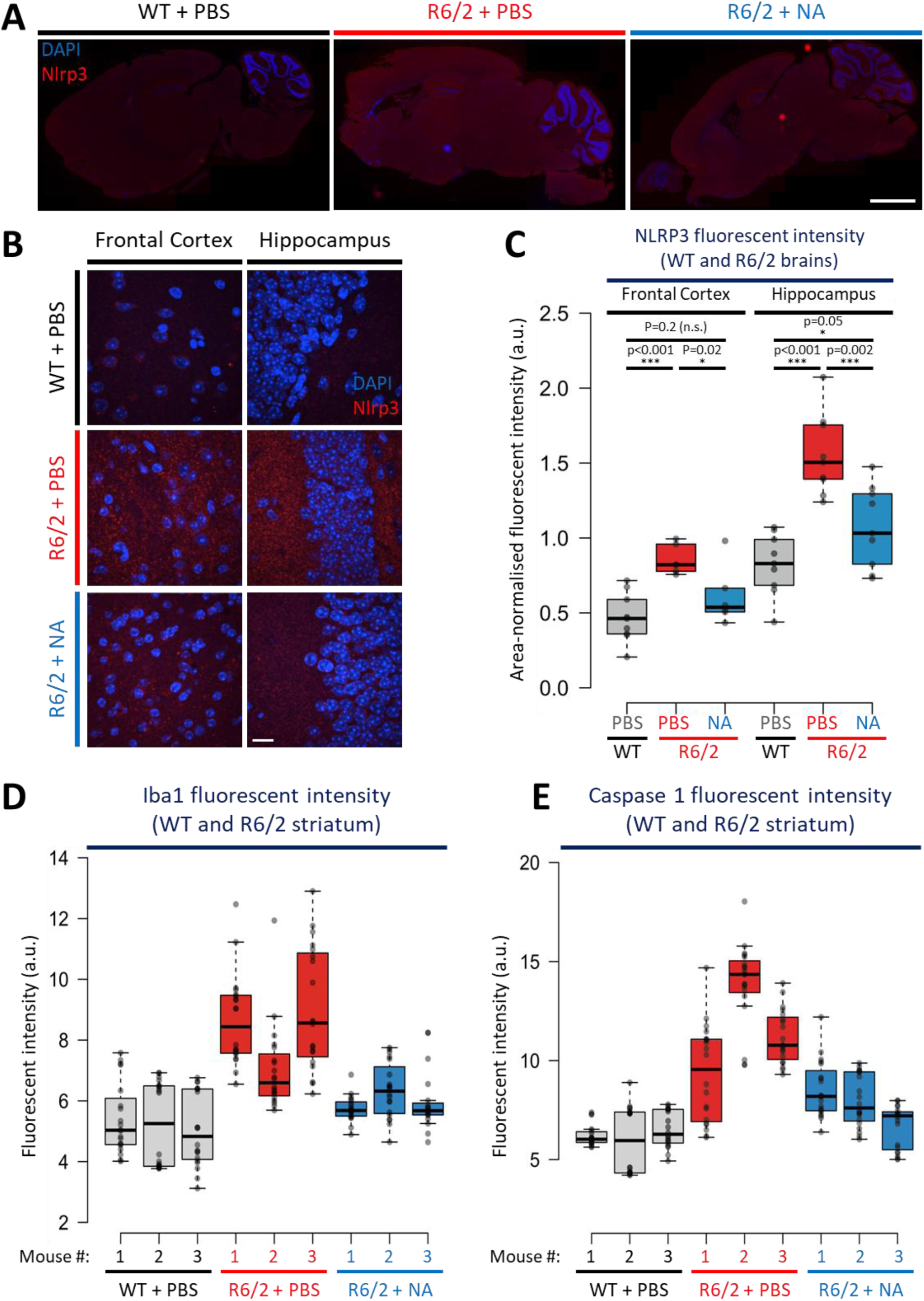
NA diminished neuroinflammation and pyroptosis related pathways in R6/2 striata. A-B) Representative A) fluorescent lightsheet images outlining Nlrp3 staining in whole brain slices and B) 40x confocal images outlining Nlrp3 staining in the striatum, frontal cortex, and cerebellum, of 10-week-old vehicle-treated WT mice and R6/2 mice +/−NA. Red signal denotes Nlrp3 signal and blue signal denotes DAPI-stained nuclei. Scale bar = 2000 μm (A) and 16 μm (B). **C)** quantifications of area-normalised Nlrp3 fluorescent intensity (a.u.) per whole frontal cortex or hippocampus intensity. **D-E)** Individual mouse quantifications Iba1 and Caspase 1 signal intensity per 40x image in the striatum of 10-week-old R6/2 mice +/−NA (n = 10 images per mouse with n = 3 mice per condition. For all box plots: box denotes 1^st^ to 3^rd^ quartiles, black lines denote median, Tukey whisker extents.

**Extended Figure 6.**
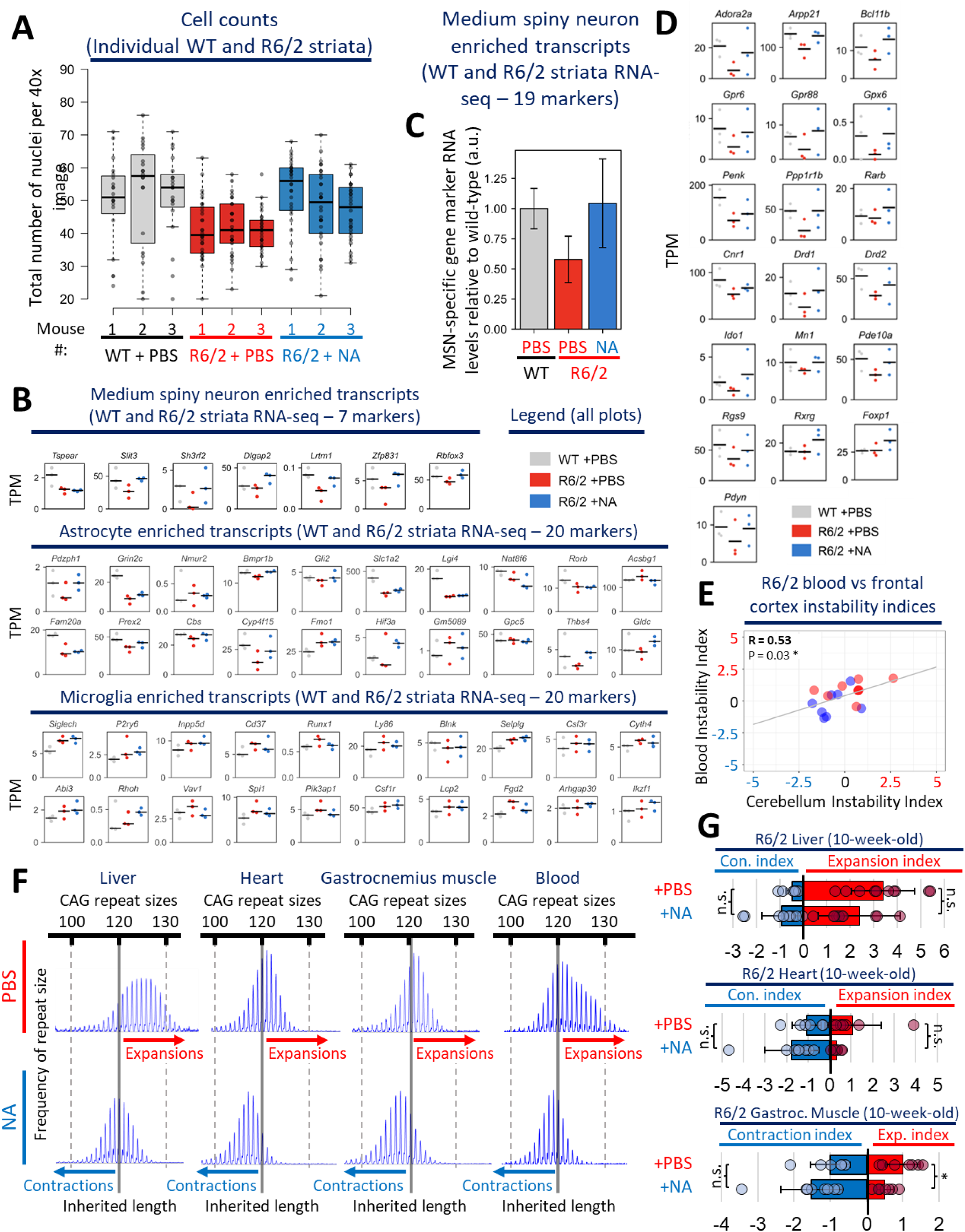
NA prevented striatal cell depletion and modulated instability in peripheral tissues. **A)** Individual mouse quantification of the number of striatal cells in 10-week-old vehicle-treated WT mice and R6/2 mice +/−NA (n = ∼3000-4500 total cells counted over 20-30 40x images per mouse per condition, 2-3 technical replicates per mouse, and 3 mice per condition). Box denotes 1^st^ to 3^rd^ quartiles, black lines denote median, Tukey whisker extents. **B)** Expression of individual markers used for striatal cell type composition quantifications of medium spiny neuron (7 markers), astrocytes (20 markers), and microglia (20 markers) in 10-week-old vehicle-treated WT and R6/2 HD mice +/−NA striata (n = 3 mice per condition). **C)** Striatal cell type composition quantifications of n = 19 medium spiny neuron (MSN) cell-type specific markers expression levels derived from 10-week-old vehicle-treated WT and R6/2 HD mice +/−NA striata (n = 3 mice per condition). **D)** Expression of individual 19 markers used for striatal cell type composition quantifications of medium spiny neurons in 10-week-old vehicle-treated WT and R6/2 HD mice +/− NA striata (n = 3 mice per condition). **E)** Pearson correlations comparing blood instability indices to cerebellar instability indices, * denotes p<0.05. **F)** Representative fragment length analysis (FLA) scans from Peak Scanner software (ThermoFisher Scientific) outlining liver, heart, muscle, and blood repeat instability in 10-week-old R6/2 mice +/−NA. Inherited length (120 CAG) is denoted with a solid gray line, additional reference lengths (+/−20 CAG units from the 120 CAG) are denoted using dashed gray lines. Expansion-biased peaks are denoted with a red arrow while contraction-biased peaks are denoted with a blue arrow. **G)** Blood repeat expansion indices denoting the average expansion/contraction indices in the liver, heart and gastrocnemius muscle of 10-week-old R6/2 mice treated with vehicle (n = 9) or NA (n = 8). Bars denote average instability indices, error bars represent standard deviation from the mean, and circles represent values for individual mice. Statistics: one-tailed Mann-Whitney U test, * denotes p<0.05, n.s. is not significant.

## Funding

The authors declare no conflicts of interest. T.G.-D. is supported by Postdoctoral Researcher Fellowships from the Hereditary Disease Foundation and the Fox Family Foundation. C.E.P. is supported by the Canadian Institutes of Health Research (FRN148910 and FRN-173282), the Natural Sciences and Engineering Research Council of Canada (RGPIN-2016-08355 and RGPIN-2016-06355/498835), The Hereditary Disease Foundation, the Petroff Family Foundation, Tribute Communities, the Marigold Foundation, the Kazman Family Foundation, and the Fox Family Foundation. C.E.P. holds a Tier 1 Canada Research Chair in Disease-Associated Genome Instability. J.S.S. is supported by the Center for Cancer Research of the National Cancer Institute, NIH (Z01-BC011585 09). E.T.W. is supported by the National Institute of Health (R01-AG058636).

## Acknowledgements

TGD, IQ, ST, MM conducted FLA and IF experiments. SYK, TGD conducted mouse phenotyping experiments and data processing; KF, KY, SCL, TKP, PW maintained mouse colonies and conducted surgical pump implantation. MK conducted correlation analyses. MN conducted RNA-seq; CPK, MK, TGD processed and analysed RNA-seq data. AC, AT, LMB conducted NfL analysis. CRH, JY provided technical support. TGD, GBP, JP, KN, LMB, PW, JSSJr, MK, PWF, ET, and CEP provided intellectual support. TGD and CEP planned experiments. TGD, SYK, IQ, CPK, and CEP wrote significant portions of the manuscript, and all authors provided edits.

## Notes

### Competing Interest Statement

CE Pearson, K Nakatani, & M Nakamori share a patent on the use of Napthyiridine Azaquinolone.

